# NucVerse3D: Generalizable 3D nuclear instance segmentation across heterogeneous microscopy modalities

**DOI:** 10.64898/2026.02.05.704108

**Authors:** Jorge Vergara, Cristian Perez-Gallardo, Ricardo Velasco, Dilan Martinez, Diego Badilla, Esteban G. Contreras, Pamela Guevara, Fabián Segovia-Miranda, Hernán Morales-Navarrete

## Abstract

Accurate three-dimensional (3D) nuclear instance segmentation is a prerequisite for quantitative phenotyping in volumetric microscopy, yet remains challenging in densely packed tissues, irregular nuclear morphologies, and across heterogeneous imaging modalities. Here we present NucVerse3D, a deep-learning framework for generalized 3D nuclei instance segmentation that combines a residual attention 3D U-Net architecture with a reversible gradient-field representation for robust centroid-aware instance reconstruction. NucVerse3D is trained end to end in 3D using modality-agnostic preprocessing and isotropic scale normalization, enabling deployment across confocal microscopy, two-photon microscopy, light-sheet microscopy, micro–computed tomography, and scanning electron microscopy volumes. We benchmarked NucVerse3D on seven volumetric datasets spanning multiple species and tissues, comprising more than forty thousand manually annotated nuclei, including newly released ground-truth datasets of mouse liver tissue (control and hepatocellular carcinoma) and *Drosophila* brain glial nuclei. Across datasets, NucVerse3D achieved consistently high precision, recall, F1-score, and average precision, and outperformed the state-of-the-art methods particularly in dense and irregular settings, while remaining competitive on simpler cases. A single generalized model trained on pooled data matched the performance of dataset-specific models, and ablation experiments demonstrated that preprocessing and scale normalization substantially contribute to performance under strict intersection-over-union criteria. To demonstrate the biomedical utility of NucVerse3D, we applied it to three-dimensional liver images from a mouse model of hepatocellular carcinoma (HCC). High-fidelity, nucleus-by-nucleus segmentation enabled the quantification of the Nuclear Decoupling Score (NDS), which captures deviations in nuclear DNA–volume coupling at the single-nucleus level. NDS analysis revealed a progressive increase in nuclear abnormalities within tumor regions, forming spatially coherent domains of dysregulated nuclei and highlighting NDS as a potential quantitative biomarker of dysplastic and tumor tissue. Together, NucVerse3D provides a robust and generalizable solution for 3D nuclear instance segmentation and enables quantitative nuclear phenotyping across imaging modalities.

**Highlights:**

- NucVerse3D provides accurate 3D nuclear instance segmentation across modalities
- Residual attention and gradient fields enable robust separation of dense nuclei
- New 3D annotated datasets of mouse liver and *Drosophila* brain are released
- A generalized model achieves performance comparable to dataset-specific training
- 3D nuclear phenotyping reveals spatially organized nuclear abnormalities in HCC

## Introduction

Three-dimensional (3D) nuclei segmentation represents a fundamental challenge in computational pathology and developmental biology, serving as the critical first step for quantitative analysis of cellular morphology, tissue architecture, and disease progression(Caicedo et al., 2019b; Falk et al., 2019; Martínez-Torres et al., 2024). Advances in volumetric microscopy techniques - including light-sheet fluorescence microscopy (LSFM) also known as selective plane illumination microscopy (SPIM)(Huisken et al., 2004; Power and Huisken, 2017), confocal(Pawley, 2006), and multiphoton imaging(Helmchen and Denk, 2005) - have enabled unprecedented visualization of nuclear dynamics in intact tissues(Keller et al., 2008; Tomer et al., 2012). However, the increasing complexity and scale of these 3D datasets have exposed significant limitations in current computational approaches for nuclear segmentation, particularly when analyzing densely packed, irregularly shaped nuclei in tissues such as developing embryos, tissues with high cellular density (e.g. liver), or pathological conditions (e.g. tumors)(Hollandi et al., 2022; Mougeot et al., 2022).

Precise 3D segmentation of densely packed nuclei is essential for understanding complex biological systems, such as developmental and pathophysiological studies where nuclear organization reflects spatial patterning and cell lineage relationships(Amat et al., 2014; Martínez-Torres et al., 2024; McDole et al., 2018). Accurate reconstruction of nuclear geometry and positioning in such contexts requires segmentation methods capable of resolving thousands of irregularly shaped nuclei within tightly confined tissue volumes. Similarly, in cancer research, robust tools for nuclear segmentation are critical, as nuclear morphology serves as a key diagnostic and prognostic biomarker(Abel et al., 2024; Fischer, 2020; Zink et al., 2004). In fact, nuclear size, shape, and spatial arrangement are well-established morphological biomarkers in hepatocellular carcinoma (HCC), with strong correlations to tumor grade and clinical outcomes(Gheonea et al., 2014; Santos et al., 2022). However, most evidence supporting these features comes from classical 2D assessments, such as hematoxylin–eosin or immunofluorescence staining of thin tissue sections, which capture only a subset of the underlying nuclear complexity. In contrast, only full 3D analysis allows for a reliable quantification of properties such as nuclearity (number of nuclei per cell) and ploidy (total amount of DNA in a cell nucleus), features that are particularly important in polyploid tissues like the liver (Chao et al., 2017; Wang et al., 2021; Wilson and Duncan, 2023). These applications place stringent demands on segmentation algorithms, which must be able to handle: (1) high nuclear densities, (2) substantial morphological variability, from spherical to highly pleomorphic nuclei, and (3) a range of imaging modalities with differing resolutions and signal-to-noise characteristics.

Current methods for 3D nuclei segmentation can be broadly categorized into three approaches: (1) intensity thresholding followed by morphological operations(Morales-Navarrete et al., 2015; Roeder et al., 2012), (2) classical machine learning using hand-crafted features(Arganda-Carreras et al., 2017; Berg et al., 2019), and (3) deep learning-based methods(Ali et al., 2025; Liu et al., 2021). While deep learning approaches - particularly those based on U-Net architectures(Bettancourt et al., 2024; Ronneberger et al., 2015; Weigert et al., 2018) - have demonstrated superior performance, most existing solutions suffer from several limitations when applied to complex biological samples. Many methods rely on assumptions of convex or slightly ellipsoidal nuclear shapes, which frequently fail for irregular or non-spherical nuclei commonly observed in both physiological cell types (e.g., endothelial cells, fibroblasts, muscle cells) and pathological conditions(Singh and Lele, 2022; Skinner and Johnson, 2017; Stringer et al., 2021; Weigert et al., 2019). Moreover, some 3D implementations are essentially 2D models extended to volumetric data through patch-based processing or z-stack projection, resulting in loss of 3D contextual information(Caicedo et al., 2019a; Weigert et al., 2019; Zhou et al., 2025). Moreover, many commonly used benchmarks for training are based on 2D images with primarily isolated or sparsely distributed nuclei, offering limited relevance for evaluating performance in 3D, densely populated, and heterogeneous tissues, scenarios where key challenges such as light distortion caused by the point spread function (PSF) are not represented(Greenwald et al., 2022; Mahbod et al., 2024; Stringer et al., 2021; Zhou et al., 2025).

To address these challenges, we present NucVerse3D, a novel deep learning framework specifically designed for accurate 3D segmentation of complex nuclear architectures. Our approach combines three key innovations: A 3D Residual Attention U-Net architecture that integrates residual blocks with cross-dimensional attention gates to simultaneously capture fine-scale nuclear boundaries and global contextual information(He et al., 2015; Ni et al., 2019; Oktay et al., 2018). A hybrid output representation that combines binary segmentation with centroid prediction, enabling robust instance separation in post-processing. End-to-end 3D training on a comprehensive collection of datasets spanning multiple tissue types (liver, brain, whole-body), a range of species (mouse, *Drosophila*, zebrafish, C. elegans), and various microscopy modalities (confocal, two-photon, light-sheet, microCT, and scanning electron microscopy). We further contribute three new 3D benchmark datasets: normal mouse liver tissue, mouse liver tissue with HCC and *Drosophila* brain glia nuclei, that specifically target challenging cases of nuclear crowding and morphological heterogeneity. These datasets are publicly available, filling an important gap in current community resources.

Through extensive validation on 7 datasets comprising >40,000 3D annotated nuclei, we demonstrate that NucVerse3D outperforms existing state-of-the-art methods for dense, irregular nuclei while maintaining comparable performance on simpler cases. To demonstrate the practical utility of NucVerse3D in a biologically relevant context, we applied it to analyze liver tumors from a well-established mouse model of HCC. We further demonstrated its translational relevance by applying it to volumetric liver imaging from an HCC mouse model, where accurate instance segmentation enables downstream 3D phenotyping at single nuclei scale. Using a control-derived DNA content vs nuclear volume model, we detected nuclear DNA content–volumne decoupling that increases from normal tissue to tumor-adjacent tissue to tumor tissue regions and forms spatially coherent domains of abnormal nuclei, suggesting localized regions of nuclear dysfunction and suggesting a potential biomarker for liver pathology. These findings highlight the potential of our approach for uncovering subtle yet biologically significant alterations in nuclear architecture associated with liver tumor progression.

## Results

### NucVerse3D: an end-to-end framework for robust 3D nuclear instance segmentation

Current 3D nuclear segmentation methods face fundamental limitations when applied to densely packed, morphologically heterogeneous tissues and to datasets acquired across diverse imaging modalities. Many existing approaches rely on simplifying assumptions, such as near-convex nuclear shapes(Weigert et al., 2019) or high signal-to-noise conditions, that rarely hold in real biological imaging scenarios. Others extend 2D architectures to volumetric data without explicitly modeling 3D nuclear geometry(Stringer et al., 2021), resulting in incomplete separation of touching nuclei, specially in the axial direction. These constraints highlight the need for a method that is both modality-agnostic and shape-agnostic, and capable of delivering accurate 3D segmentation across different normal, and pathological tissues.

NucVerse3D addresses these gaps through an integrated, end-to-end framework designed specifically for volumetric nuclear segmentation at scale (Fig. 1). The pipeline begins with raw 3D z-stacks, which undergo a modality-agnostic preprocessing stage that performs illumination correction, alignment, unsupervised deconvolution, and intensity normalization (Fig. 1A). This step harmonizes image characteristics across microscopes, reducing variability induced by optical distortions, depth-dependent attenuation, and acquisition settings. At the core of the pipeline is a residual attention U-Net 3D architecture that simultaneously predicts two complementary outputs: a 3D probability mask and a 3D gradient field (Fig. 1B). Whereas the probability mask provides voxel-level nuclear likelihoods, the gradient field encodes directional information toward nuclear centroids along all three spatial dimensions. By learning this geometric structure directly from data, the model builds an internal representation of nuclear topology that is not constrained by spherical or star-convex shape assumptions.

**Figure 1:**
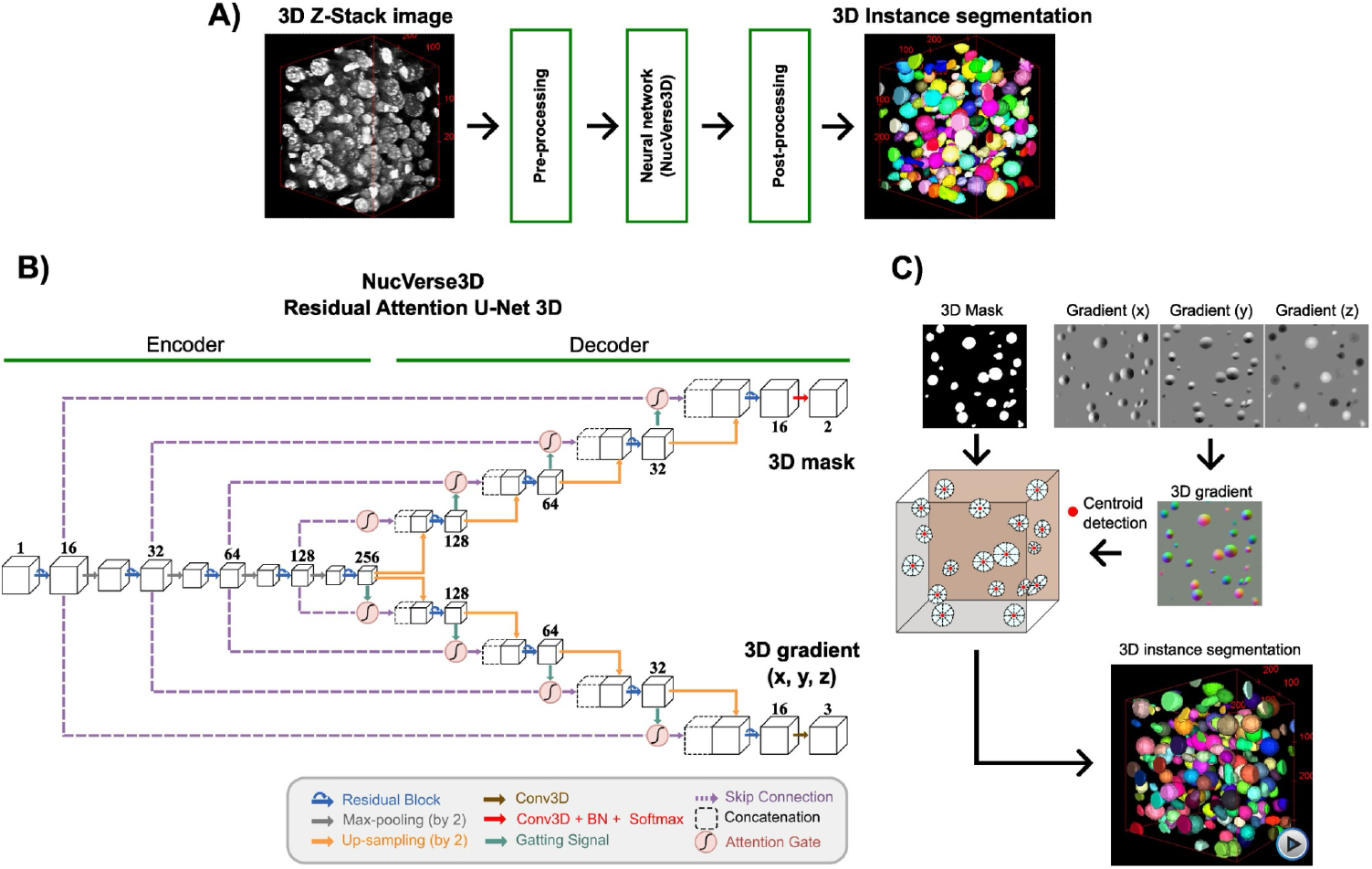
NucVerse3D pipeline and model for 3D nuclear instance segmentation. **(A)** End-to-end segmentation workflow. A raw 3D Z-stack image containing nuclei is first subjected to preprocessing (e.g., intensity normalization and unsupervised deconvolution). The preprocessed volume is then processed by the NucVerse3D neural network, which operates directly in 3D. Finally, a post-processing step converts the network outputs into a labeled 3D instance segmentation, in which each nucleus is assigned a unique identity and color. **(B)** Architecture of the NucVerse3D Residual Attention 3D U-Net, consisting of an encoder– decoder structure with residual blocks, skip connections, and attention gates. The network jointly predicts a 3D nuclear mask and 3D gradient maps (x, y, z). **(C)** Instance separation strategy. Predicted nuclear masks and gradient fields are combined to detect nuclear centroids and separate touching nuclei, yielding the final 3D nuclear instance segmentation (Supplementary Movie 1).

During post-processing, these outputs are combined to derive final nuclear instances (Fig. 1C). Local attractors within the predicted gradient field identify candidate centroids, while the probability mask provides spatial support for voxel assignment. A gradient-descent–based grouping mechanism then aggregates voxels that converge to the same attractor, reconstructing each nuclear volume with high boundary fidelity. This data-driven strategy replaces traditional watershed or distance-transform heuristics, leading to robust separation of tightly packed or irregularly shaped nuclei (see Supp. Movie 1).

Together, these components form a fully reversible, end-to-end 3D instance segmentation framework that is inherently adaptable to variations in nuclear morphology, imaging contrast, voxel anisotropy, and signal-to-noise characteristics. NucVerse3D thereby overcomes key limitations of existing methods, providing a unified, generalizable solution for large-scale 3D biomedical image analysis across species, tissues, and imaging technologies.

### Comprehensive cross-modality benchmarking and comparative performance analysis

A major limitation for 3D nuclear segmentation is the scarcity of high-quality, manually annotated volumetric datasets. While many neural networks rely on abundant 2D annotations, high-quality 3D ground truth remains rare due to the complexity of volumetric manual annotation. To address this gap, we generated and publicly released two new manually annotated 3D datasets: (i) Normal mouse liver and (ii) *Drosophila* Brain Glia (Supp. Table 1). Together, they contain more than 5000 fully annotated nuclei and were explicitly designed to represent the morphological challenges most segmentation algorithms fail to capture. Whereas, the mouse liver dataset includes large hepatocyte nuclei, irregular polyploid nuclei, and small elongated nuclei from non-parenchymal cells, many of which are touching or partially overlapping, conditions that frequently cause over-merging in conventional pipelines; the *Drosophila* glial dataset features extreme size heterogeneity within the same volume, reflecting the biological diversity of glial subtypes. These datasets provide a needed benchmark for evaluating segmentation in highly crowded, morphologically heterogeneous tissues and are publicly available to support further community development.

To rigorously evaluate the generalization capability of NucVerse3D, we benchmarked its performance across six heterogeneous 3D microscopy datasets spanning a wide range of voxel resolutions, nuclear morphologies, and imaging modalities, including (Supp. Table 1 and Figure 2): *C. elegans* embryos (confocal), characterized by small, compact nuclei at high density(Hirsch and Kainmueller, 2020; Long et al., 2009); whole mouse brain (mesoSPIM), containing very large nuclei (mean volume ∼29800 µm^3^) at relatively low density(Achard et al., 2025); mouse visual cortex (microCT), with intermediate-sized, nearly spherical nuclei and moderate density(Lin et al., 2021); zebrafish brain (SEM), presenting high-density nuclei (∼1.0×10^6^ nuclei/mm^3^) with low sphericity(Lin et al., 2021); *Drosophila* brain (confocal), with pronounced variability in shape and heterogeneous nuclear staining, and adult mouse liver (two-photon), containing both spherical and non-spherical nuclei and substantial crowding by non-parenchymal cells. Together, these datasets capture a wide spectrum of nuclear configurations encountered in developmental, anatomical, and pathological imaging: from small and nearly spherical (*C. elegans*) to highly pleomorphic (liver), from low-density to extremely crowded (SEM zebrafish), and from high-resolution confocal volumes to coarse, mesoscopic light-sheet acquisitions (Supp. Table 1 and Figure 2).

**Figure 2:**
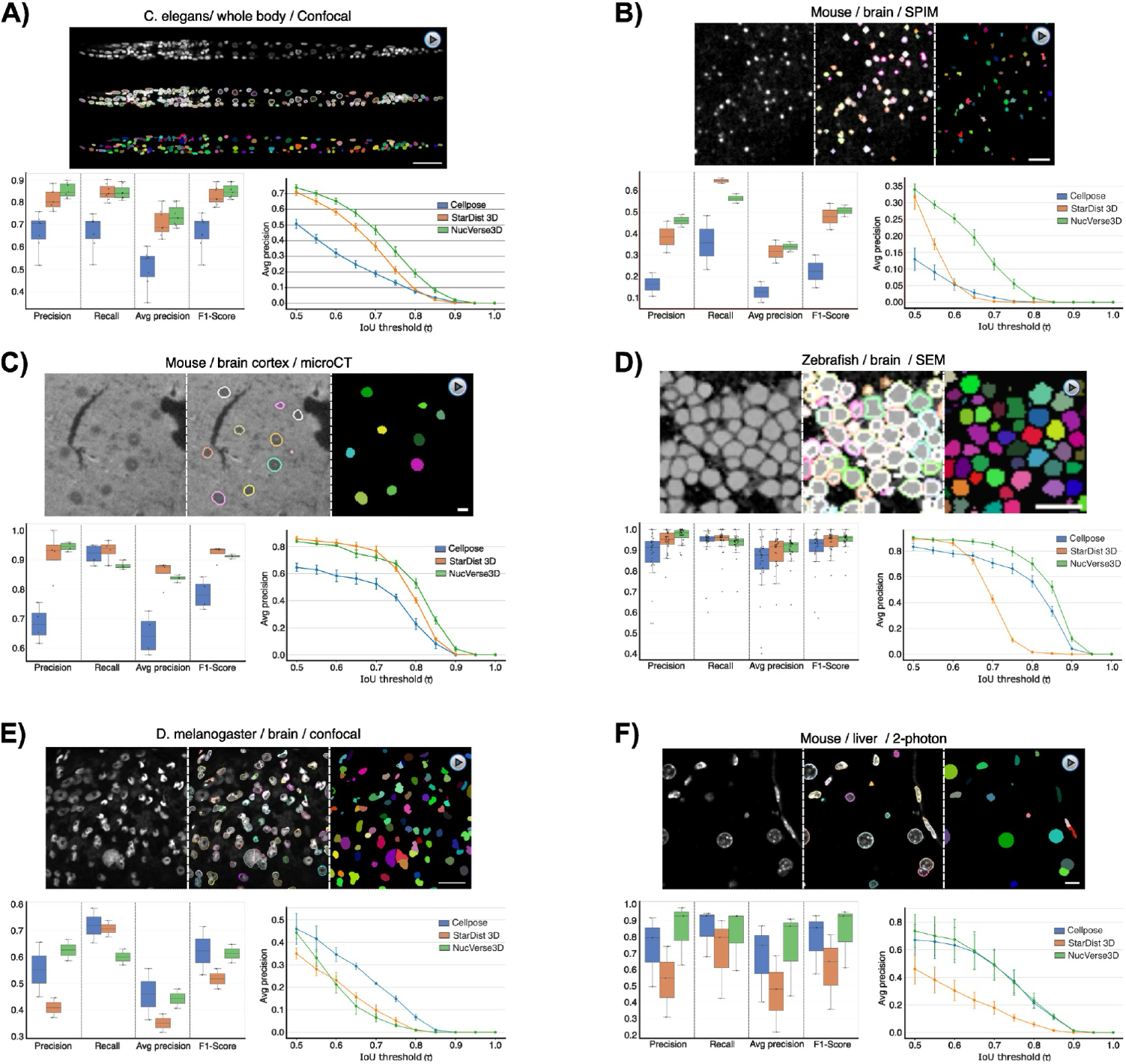
Cross-dataset evaluation of 3D nuclei segmentation methods. Qualitative and quantitative evaluation of 3D nuclear instance segmentation across six volumetric microscopy datasets, comparing NucVerse3D with Cellpose and StarDist 3D. **(A)** *C. elegans* whole-body imaging by confocal microscopy. **(B)** Mouse brain imaged by light-sheet mesoSPIM. **(C)** Mouse visual cortex acquired by microCT. **(D)** Zebrafish brain imaged by scanning electron microscopy (SEM). **(E)** *Drosophila melanogaster* brain imaged by confocal microscopy. **(F)** Adult mouse liver imaged by two-photon microscopy. For each dataset, representative images show the raw input image, the predicted nuclei displayed as outlines, and the predicted nuclei shown as instance masks (colored). The bottom panels report quantitative performance metrics (precision, recall, average precision, and F1-score), where each point represents a test volume; boxes indicate the interquartile range and median. In addition, average precision curves are shown, evaluated at the instance level. For each dataset, an associated 3D visualization of the segmentation results is provided in Supplementary Movies 2–7. Error bars represent mean ± standard error of mean across test volumes (*C. elegans* confocal, n = 7; mouse brain SPIM, n = 2; mouse brain microCT, n = 4; zebrafish brain SEM, n = 27; *Drosophila* brain confocal, n = 2; mouse liver 2-photon, n = 3).

When benchmarked against two widely used 3D instance-segmentation baselines, StarDist 3D(Weigert et al., 2019) and Cellpose 3D(Stringer et al., 2021), NucVerse3D achieved consistently strong and often superior performance across a diverse set of imaging modalities and biological systems (Fig. 2A–F). Across fluorescence-based datasets, NucVerse3D typically yielded high F1-scores and balanced precision–recall trade-offs, indicating reliable detection of nuclei while maintaining accurate instance separation. Importantly, improvements were not limited to a single dataset or modality: NucVerse3D remained competitive in confocal whole-body C. elegans imaging (Fig. 2A), SPIM whole-brain data (Fig. 2B), confocal *Drosophila* brain tissue (Fig. 2E), and two-photon mouse liver volumes (Fig. 2F), demonstrating broad generalization under heterogeneous resolution, contrast, and nuclear packing regimes.

Visual comparisons (Fig. 2A–F, Sup. movies 2-7) show that NucVerse3D consistently resolves nuclei with high fidelity, even under strong imaging challenges such as dense nuclear packing, anisotropic blur, pleomorphic morphologies, low signal-to-noise ratios, and modality-specific artifacts. In contrast, baseline methods frequently exhibited over-segmentation in crowded regions or failed to separate irregularly shaped nuclei. Quantitative analyses corroborate these observations. NucVerse3D provides the highest precision, recall, average precision (AP), and F1-scores across all six datasets. Across species and modalities, NucVerse3D maintains Precision and Recall typically over 0.80, F1-scores also mostly over 0.80 and Average precision (AP) at IoU = 0.5 above 0.7 for most of the datasets, often exceeding 0.85. Its performance remains stable even in challenging contexts, including datasets with non-spherical or pleomorphic nuclei, such as liver and *Drosophila* brain tissues, where nuclear boundaries are highly irregular and densely packed. Altogether, these results demonstrate that NucVerse3D achieves state-of-the-art segmentation accuracy across species, tissues, imaging modalities, and nuclear morphologies, providing a generalizable solution for large-scale 3D biomedical image analysis and filling a critical gap in current segmentation resources through the release of new, challenging 3D ground-truth datasets.

A key differentiator emerges when performance is evaluated under increasingly strict overlap criteria. Beyond aggregate metrics at a single threshold, Fig. 2 reports average precision as a function of the IoU threshold τ, providing a stringent assessment of boundary fidelity and instance-level correctness. In multiple datasets, competing methods exhibited a pronounced deterioration in AP as τ increased, consistent with boundary imprecision, instance merging, or over-segmentation errors becoming increasingly penalized. In contrast, NucVerse3D showed substantially higher AP retention at high IoU thresholds, indicating that it preserves instance geometry and boundary accuracy more reliably. This effect is particularly evident in datasets where precise 3D delineation is challenging due to dense packing, anisotropy, or reduced SNR (e.g., SPIM brain volumes and two-photon liver tissue; Fig. 2B,F). Notably, for modalities that depart from conventional fluorescence microscopy, such as microCT (Fig. 2C) and SEM (Fig. 2D), NucVerse3D remained robust, maintaining strong AP across a broad IoU range and avoiding the sharp high-τ collapse observed for at least one baseline in these settings. Together, these trends suggest that NucVerse3D’s instance reconstruction is not only accurate in terms of detection but also more faithful in 3D morphology, a property that is critical for downstream quantitative phenotyping where small boundary errors can propagate into biased volumetric or intensity-derived measurements.

Overall, the benchmark results demonstrate that NucVerse3D provides more stable cross-dataset behavior than current state-of-the-art approaches, with performance advantages that become most apparent under strict evaluation regimes that emphasize true instance quality rather than coarse overlap. This stability across species, tissues, and modalities is particularly relevant for large-scale 3D analysis pipelines, where robustness to imaging variability and morphological heterogeneity is essential for reproducible quantification and biological interpretation.

### Performance of a generalized model trained across all datasets

A recurring practical limitation of deep-learning segmentation pipelines is the need to train and maintain dataset-or modality-specific models, which reduces reproducibility and complicates deployment in heterogeneous imaging studies. To test whether NucVerse3D can instead operate as a single general-purpose model without sacrificing accuracy, we trained a generalized network on pooled data from all benchmark datasets and compared it directly against dataset-specific models under controlled training conditions. To avoid confounding effects due to dataset imbalance, we standardized the training volume per dataset, randomly sampling 1,000 patches per dataset (800 training, 200 validation) for both the generalized and the specialized models. In addition, all volumes were isotropically rescaled so that the average nuclear radius was approximately matched across datasets (Methods), minimizing trivial scale-driven domain gaps while preserving modality-specific appearance characteristics.

Across six imaging modalities and biological systems, the generalized model achieved comparable performance to specialized models across all core metrics (Fig. 3A). Precision and recall remained high for most datasets, and critically, the generalized model preserved F1-score and average precision at levels closely tracking the dataset-specific baselines for most datasets. This indicates that pooling heterogeneous data does not dilute performance, but instead enables the network to learn shared, transferable priors for nuclear geometry and separation. Importantly, the agreement between generalized and specialized models extends across large differences in voxel size and nuclear morphology, suggesting that the combination of 3D residual-attention features and the gradient-field instance representation yields a robust, modality-agnostic segmentation strategy.

**Figure 3:**
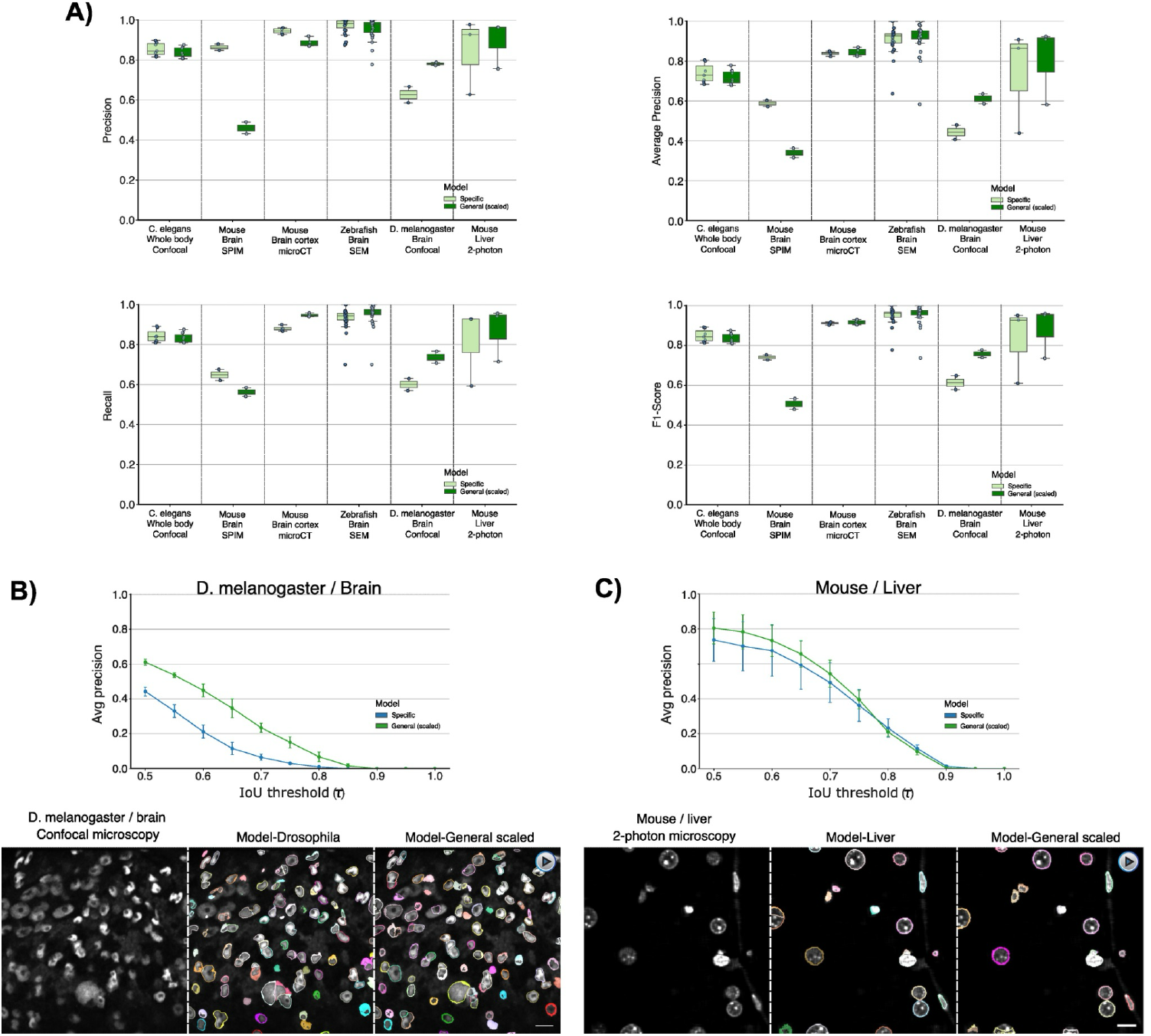
General vs. Specific Model Comparison. Instance-level segmentation performance of dataset-specific and generalized (scaled) NucVerse3D models across multiple volumetric microscopy datasets. **(A)** Box plots of precision, recall, average precision, and F1-score across six datasets. Each point represents a test volume; boxes indicate the interquartile range and median. **(B)** Average precision curves for the *D. melanogaster* brain and mouse liver **(C)** datasets. Representative qualitative examples show raw images, and predicted nuclear outlines for dataset-specific and generalized models (Supplementary Movies 8-9). Error bars represent mean ± standard error of mean across test volumes (*C. elegans* confocal, n = 7; mouse brain SPIM, n = 2; mouse brain microCT, n = 4; zebrafish brain SEM, n = 27; *Drosophila* brain confocal, n = 2; mouse liver 2-photon, n = 3).

The IoU-threshold analysis further supports this conclusion. In both *Drosophila* brain confocal and mouse liver two-photon datasets (two regimes that strongly challenge instance separation due to heterogeneous nuclear sizes and shapes), the generalized model maintains an average precision–IoU curve closely aligned with the specialized model or even improves in the case of the *Drosophila* (Fig. 3B–C; Supp. Movie 8). Similar behaviour is observed for the other dataset except Mouse brain where the specialized model performs better than the general one (Supp. Fig. 1) While AP decreases as the IoU threshold becomes more stringent, as expected for any instance segmentation method, the generalized model shows a smooth, gradual decay rather than a sharp collapse, indicating that its performance is not driven by coarse overlap alone but retains meaningful boundary fidelity even under strict criteria. The qualitative comparisons reinforce these quantitative trends: in both datasets, the generalized model reconstructs nuclear instances with boundary adherence and separation quality comparable to dataset-specific training, including in crowded regions where over-merging or fragmentation typically occurs.

Together, these results demonstrate that NucVerse3D can be deployed as a single unified segmentation model trained across heterogeneous datasets, achieving accuracy comparable to specialized models while dramatically reducing the burden of modality-specific retraining. This property is particularly relevant for large-scale 3D microscopy workflows, where datasets frequently span multiple microscopes, staining protocols, tissues, and spatial resolutions, and where robust generalization is essential for reproducible, scalable quantitative biology.

### Ablation analysis reveals the importance of image scaling and preprocessing

To quantify the contribution of key pipeline components to overall segmentation performance, we performed an ablation analysis in which we removed (i) the image scaling step used to harmonize object size across datasets and (ii) the preprocessing module comprising illumination correction, unsupervised deconvolution, and intensity normalization. The resulting reduced models were evaluated on all benchmark datasets using the same metrics as in Figure 3, including precision, recall, F1-score and average precision across IoU thresholds.

Removing either component led to a consistent degradation in performance relative to the complete NucVerse3D pipeline (Fig. 4A–F), confirming that high-quality 3D instance segmentation is not achieved by network design alone but emerges from the joint optimization of data standardization and model inference. The impact of scaling was particularly pronounced and dataset-dependent. In several datasets, most notably mouse brain cortex (microCT, Fig.4C) and *Drosophila* brain (confocal, Fig.4E), disabling scaling produced a marked reduction in precision and F1-score, accompanied by a collapse in average precision at stricter IoU thresholds, indicating an increased rate of instance fragmentation and boundary inaccuracy. In contrast, datasets with more homogeneous object scale, such as *C. elegans* confocal (Supp. Table 1), showed a smaller reduction, consistent with scaling primarily acting as a cross-domain normalization mechanism that stabilizes the geometric representation learned by the network. Only in one dataset (Mouse brain, SPIM), removing the scaling improved the segmentation quality.

**Figure 4:**
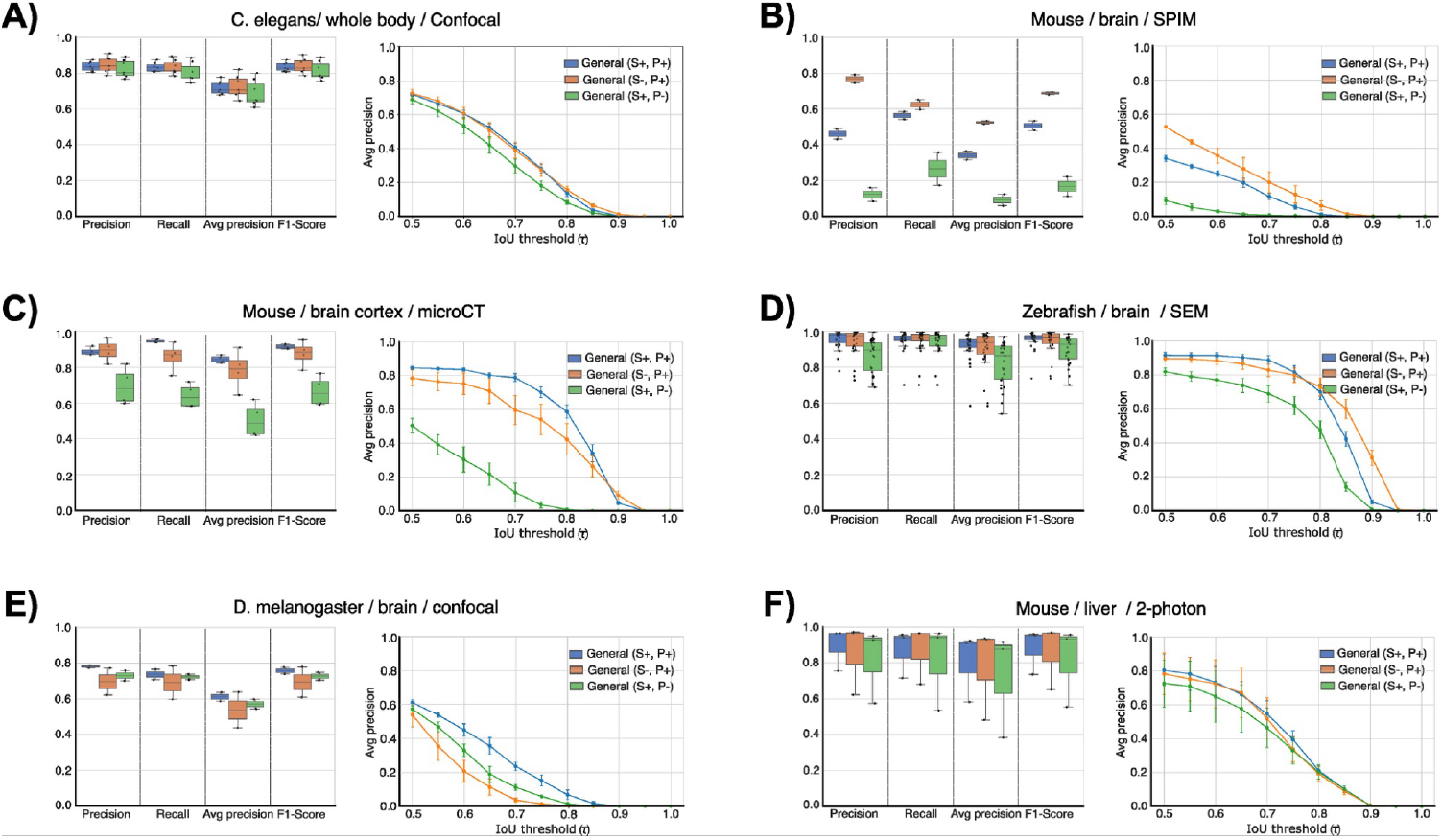
Ablation analysis of NucVerse3D pipeline components. Segmentation performance across all datasets for the full NucVerse3D model (S+, P+; blue), the model without isotropic image scaling (S-, P+; orange), and the model without preprocessing (S+, P-; green). Datasets include: **(A)** *C. elegans* (confocal), **(B)** mouse brain (SPIM), **(C)** mouse brain cortex (microCT), **(D)** zebrafish brain (SEM), **(E)** *D. melanogaster* brain (confocal), **(F)** mouse liver (two-photon). Box plots of precision, recall, average precision, and F1-score across six datasets. Each point represents a test volume; boxes indicate the interquartile range and median. Average precision curves with error bars showing mean ± standard error of mean across imaging volumes for each dataset (*C. elegans* confocal, n = 7; mouse brain SPIM, n = 2; mouse brain microCT, n = 4; zebrafish brain SEM, n = 27; *Drosophila* brain confocal, n = 2; mouse liver 2-photon, n = 3).

Ablating preprocessing produced an even more dramatic see-saw effect across modalities. In mouse brain (SPIM, Fig.4B) and mouse brain cortex (microCT, Fig.4C), removal of preprocessing caused a severe drop across all summary metrics and resulted in near-baseline average precision curves across IoU thresholds, consistent with systematic foreground– background confusion and loss of boundary contrast. In these settings, illumination artifacts, blur, and modality-dependent noise strongly affect the raw nuclear signal. Preprocessing therefore plays a critical role in restoring contrast and preventing the network from learning spurious intensity cues. By contrast, high-contrast datasets such as zebrafish brain (SEM, Fig.4D) and mouse liver (confocal Fig.4F) displayed comparatively milder degradation, suggesting that preprocessing contributes most strongly when the nuclear signal is weak or confounded by acquisition artifacts.

Importantly, the effects of both ablations were most evident under stringent evaluation conditions. While differences in coarse metrics can appear moderate in some datasets, the average precision–IoU curves revealed that both scaling and preprocessing are key to preserving accuracy at high overlap thresholds, where small boundary errors and minor instance merging events are strongly penalized. Across most benchmarks, the complete pipeline consistently retained higher AP values at strict IoU, demonstrating superior boundary fidelity and more reliable instance separation. Together, these results indicate that isotropic scaling and preprocessing are not optional refinements but constitute essential components that enable robust cross-modality generalization and stable performance in challenging 3D imaging regimes.

### Nuclear DNA content–size decoupling reveals progressive morphological disruption from control to tumor liver tissue

The liver exhibits a distinctive nuclear architecture, as a substantial fraction of adult hepatocytes are binucleated and polyploid (i.e. having more than two complete sets of chromosomes) under physiological conditions(Donne et al., 2020; Gentric and Desdouets, 2014). Numerous studies have shown that the degree of hepatocyte ploidy is dynamically regulated and correlates with liver function as well as with the progression of hepatic diseases, including hepatocellular carcinoma (HCC)(Wang et al., 2017; Zhang et al., 2018). Importantly, nuclear ploidy is directly associated with nuclear size, making nuclear morphology a meaningful structural proxy for cellular genomic content(Morales-Navarrete et al., 2015). These features render liver tissue and HCC in particular, a biologically relevant and technically demanding system for evaluating 3D nuclear segmentation approaches.

To test the robustness of NucVerse3D in pathological liver tissue, we generated full 3D reconstructions of DAPI-stained thick liver sections imaged by multiphoton microscopy from control mice and from a metabolic dysfunction–associated steatohepatitis (MASH)–associated HCC mouse model(Fujii et al., 2013). DAPI integrated intensity has previously been shown to provide a reliable quantitative readout of nuclear DNA content, enabling the assessment of DNA-content alterations in heterogeneous or diseased tissues(Morales-Navarrete et al., 2015; Segovia-Miranda et al., 2019). We first evaluated model performance using the general NucVerse3D model and the liver model. Despite not having been exposed to this data during training, both models show good performance, however, the general model achieved higher and more stable performance than the liver-specific model across precision, recall, average precision, and F1-score (Fig. 5A), demonstrating strong generalization to tumor-associated nuclear morphologies.

**Figure 5:**
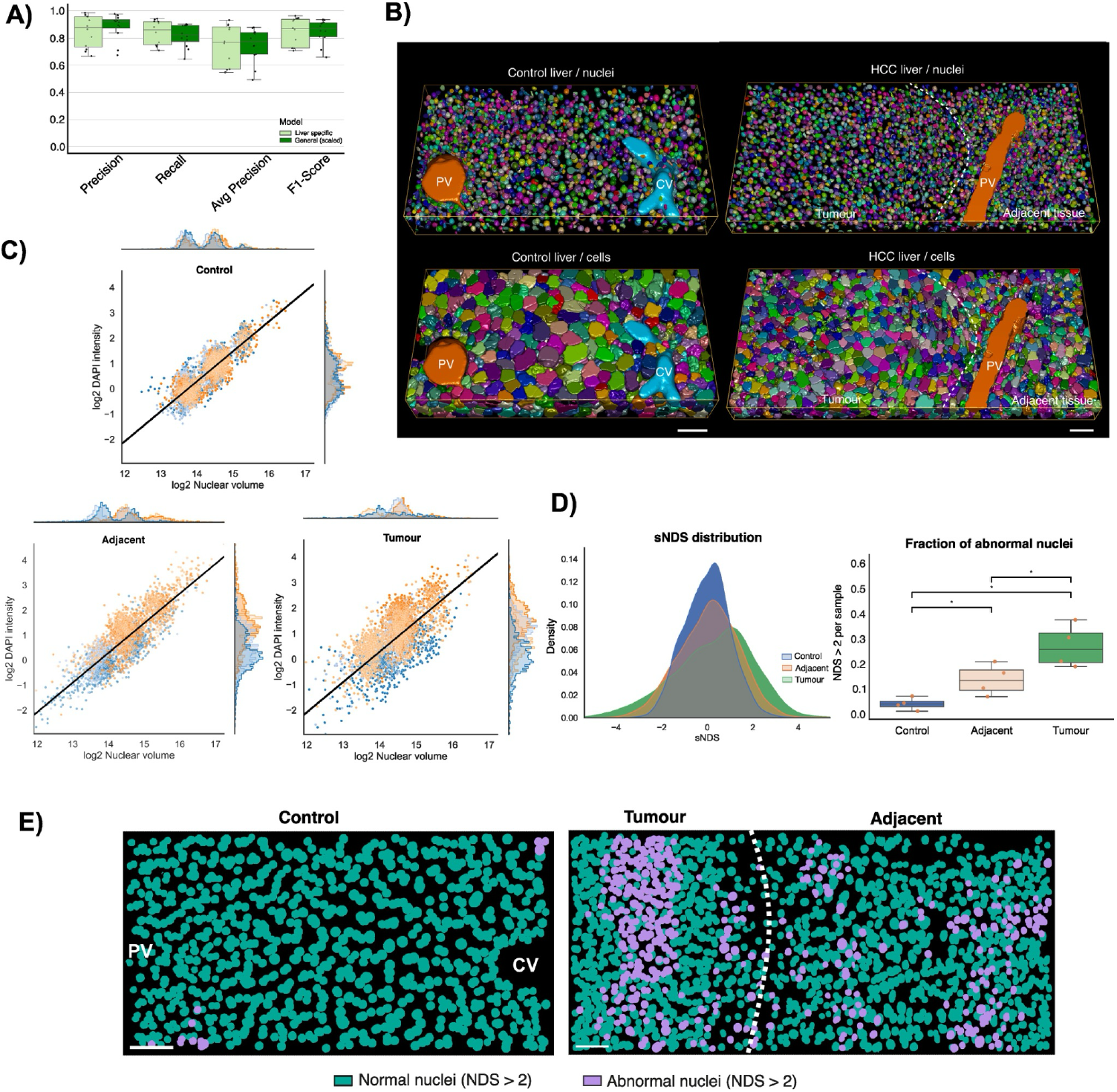
NucVerse3D reveals hidden nuclear and cellular organization patterns in HCC liver tissue. **(A)** Segmentation performance on mouse liver HCC samples for liver-specific and generalized models. (Precision, Recall, AP, F1-Score). Each point represents a sample; boxes indicate the interquartile range and median (control (n = 4) and HCC (n = 4)). **(B)** Representative 3D renderings of nuclear (top) and cellular (bottom) instance segmentations in control and HCC liver tissue. Central vein (CV), portal vein (PV), tumor, and adjacent regions are indicated. **(C)** Scatter plots of nuclear volume versus integrated DAPI intensity for control, adjacent, and tumor tissue. Each point represents a single nucleus. The solid black line indicates the control-derived scaling model, and is shown identically across all three conditions to highlight deviations from physiological coupling. **(D)** Distribution of the standardized nuclear deviation score (sNDS) across conditions (left) and fraction of nuclei classified as abnormal (sNDS > 2) per sample (right). Each point represents a sample; boxes indicate the interquartile range and median (control (n = 4) and HCC (n = 4)). Statistical comparisons were performed using one-sided Mann–Whitney U tests with Benjamini– Hochberg FDR correction (p < 0.05, p < 0.01, p < 0.001). **(E)** Maximum-intensity projection over a 50 μm depth showing the spatial distribution of nuclei classified as normal or abnormal based on sNDS in control, adjacent, and tumor regions. Scale bar 50 µm.

Using the general model, we generated complete 3D segmentations of nuclei across intact tissue, and HCC samples (Fig. 5B). General morphological analysis revealed no major differences in hepatocyte or non-parenchymal cell numbers or hepatocyte size (Supp Fig 2). Some differences were observed between healthy and HCC tissue—such as a reduction in binucleated hepatocytes, but these parameters did not distinguish tumor-adjacent tissue from tumor tissue itself (Supp. Fig 2). However, initial exploratory analyses already revealed that nuclear morphology was less regular in HCC tissue than in control liver, indicating increased heterogeneity in nuclear size (Supp. Fig 3). Consistent with this observation, analysis of the relationship between nuclear DNA content and nuclear volume showed that the tight coupling observed in control tissue progressively deteriorated in pathological regions (Supp. Fig. 4). While control nuclei exhibited a strong log–log linear correlation between integrated DAPI intensity and nuclear volume, consistently with previous studies(Morales-Navarrete et al., 2015), this correlation was partially lost in adjacent tissue and further disrupted in tumor tissue (Supp. Fig. 4). These deviations motivated the development of a quantitative framework to capture nuclear abnormalities beyond simple size measurements.

To assess whether the morphological information extracted by NucVerse3D reflects biologically meaningful alterations in nuclear physiology in samples of liver tissue with HCC, we quantified deviations between nuclear volume and DNA content using the content–volume scaling framework established from control tissue. In healthy liver, nuclear size and integrated DAPI intensity exhibited a strong log–log linear relationship with low scatter, defining a stable reference model of physiological nuclear coupling (Fig. 5C). This tight alignment reflects the well-regulated balance between nuclear ploidy and nuclear volume characteristic of hepatocytes under control conditions. In contrast, applying the same reference model to adjacent and tumor tissue revealed marked deviations: nuclei in these regions showed increased spread and systematic departures from the expected trend, indicative of disrupted DNA-content regulation and structural instability (Fig. 5C).

The distribution of signed decoupling scores (sNDS) illustrated this progressive breakdown in nuclear homeostasis. Control nuclei were narrowly centered around zero, consistent with tightly coordinated DNA content–volume coupling, while adjacent and tumor regions exhibited increasingly broadened, flattened, and shifted sNDS profiles (Fig. 5D, left). This widening reflects a substantial amplification of nucleus-to-nucleus variability in pathological tissue, aligning with known structural, genomic, and ploidy abnormalities, such as aneuploidy (i.e. abnormal number of chromosomes) in HCC(Cordier et al., 2025; Heckenbach et al., 2022). Quantifying these deviations at the sample level revealed a clear and statistically robust gradient: the fraction of nuclei classified as abnormal (NDS > 2) was lowest in control tissue, intermediate in adjacent regions, and highest in tumors (Fig 5 D-E, Supp. Fig. 5).

Projecting NDS classifications back into the 3D tissue context revealed a progressive increase in structural and spatial disorganization with disease progression. Control liver exhibited a coherent lobular architecture with predominantly normal nuclei, whereas adjacent tissue showed localized clusters of abnormal nuclei near morphologically perturbed regions (Fig 5E, Supp. Fig. 5). Tumor samples displayed the most pronounced disruption, with large contiguous domains of abnormal nuclei coinciding with architectural collapse. Together, these patterns demonstrate that nuclear content–size decoupling not only increases globally but also emerges in spatially coherent domains aligned with pathological tissue remodeling. This establishes NucVerse3D-derived nuclear phenotyping as a sensitive, spatially resolved readout of nuclear dysfunction across 3D liver tissue and highlights its potential as a quantitative biomarker of disease progression.

## Discussion

We introduce NucVerse3D, a generalizable deep-learning framework for 3D nuclear instance segmentation that integrates a residual attention architecture with a reversible gradient-field representation to achieve robust performance across diverse tissues, imaging modalities, and nuclear morphologies. Our results demonstrated that high-fidelity 3D nuclear segmentation requires more than architectural improvements alone: image scaling, image harmonization, and a biologically motivated instance-reconstruction strategy each contribute substantially to overall performance. Through extensive benchmarking and ablation experiments, we show that the full NucVerse3D pipeline consistently outperforms ablated variants, particularly in datasets with high nuclear density. Importantly, we address a key limitation in the field by releasing new manually annotated 3D datasets of mouse liver (control and HCC) and *Drosophila* brain glia, providing challenging benchmarks for community training and evaluation. Beyond methodology, application of NucVerse3D to liver tumors reveals progressive nuclear DNA content–size decoupling and spatially coherent nuclear abnormalities in hepatocellular carcinoma, highlighting the value of accurate 3D nuclear phenotyping for uncovering biologically meaningful tissue alterations.

A distinguishing feature of NucVerse3D is its ability to operate reliably across modalities ranging from confocal and two-photon microscopy to mesoSPIM light-sheet data and microCT volumes. This cross-domain stability emerges from two synergistic properties: (i) the gradient-field representation, which decouples instance separation from absolute intensity structure and is therefore less sensitive to modality-specific artifacts; and (ii) the combination of residual and attention mechanisms, which enables the model to integrate fine boundary details with wide-field contextual cues. Together, these elements provide a unified instance segmentation strategy that generalizes across species and tissues without extensive retraining. Importantly, a single model trained across heterogeneous datasets performs comparably to dataset-specific models, suggesting that NucVerse3D captures a modality-invariant geometric prior for nuclear shape and distribution. This finding is particularly relevant in the emerging landscape of 3D foundation models, where generalization beyond the training distribution remains a central challenge.

Beyond methodological advances, our work demonstrates the biological value of accurate 3D nuclear segmentation through its application to HCC, a tissue characterized by extensive alterations in binucleation and polyploidy(Wang et al., 2017; Zhang et al., 2018). Importantly, these analyses are only reliably achievable in 3D. Traditional 2D histological sections create strong artifacts, including nuclear truncation, apparent shape irregularities, and biased sampling of polyploid or binucleated cells, which confound quantitative measurements and disrupt spatial context(Garvalov and Ertürk, 2017). By quantifying nuclear DNA content–size coupling using a control-derived log–log model, we uncover a progressive and spatially coherent breakdown of nuclear homeostasis from control to adjacent and tumor tissue.This breakdown in DNA content–size coupling gives rise to a Nuclear Decoupling Score that captures subtle disruptions in nuclear homeostasis and represents a promising biomarker for next-generation 3D digital pathology(Abel et al., 2024; Hägerling et al., 2017; Tanaka et al., 2017). While conventional morphological features fail to distinguish tumor-adjacent from tumor regions, NucVerse3D reveals pronounced nuclear decoupling in tumors, often organized into spatially coherent clusters of abnormal nuclei. These pseudocluster-like structures suggest the presence of local expansions of cells sharing nuclear abnormalities, potentially reflecting clonal processes during disease progression. We hypothesize that these alterations are consistent with progressive loss of binucleation, reduction of polyploid states, and the emergence of ploidy instability such as aneuploidy, a hallmark of HCC(Cordier et al., 2025; Heckenbach et al., 2022). Although definitive confirmation will require complementary genomic validation, our results indicate that 3D image-based nuclear phenotyping can reveal spatial patterns of nuclear abnormality with strong potential as a quantitative and spatially resolved biomarker in cancer.

Despite its strengths, NucVerse3D has limitations that require further investigation. First, while the model generalizes across a wide range of microscopy modalities, performance at extremely large scales (e.g., multimeter-scale whole-organ imaging) will require further optimization of memory-efficient inference. Second, our instance reconstruction relies on gradient descent within predicted vector fields; although stable across our datasets, this process may require adaptation for tissues with extremely irregular nuclear shapes or non-intuitive topology. Third, while our biological application demonstrates quantitative and spatially resolved nuclear phenotyping, clinical translation will require integration with histopathology workflows and prospective validation in human tissue. Future work should also explore integration with 3D foundation models (e.g. SAM(Kirillov et al., 2023) or DINO(Siméoni et al., 2025)) and contrastive pretraining to further expand cross-domain generalization.

In summary, NucVerse3D provides a robust, scalable solution to one of the longstanding challenges in biological image analysis: accurate and generalizable 3D nuclear instance segmentation. By coupling methodological advances with meaningful biological validation, this work establishes a practical foundation for large-scale 3D cellular phenotyping and opens new opportunities for quantitative analysis of tissue architecture in health and disease. We anticipate that NucVerse3D will serve as a versatile tool for the imaging community and a key component in the emerging ecosystem of computational methods for high-dimensional, volumetric tissue characterization.

## Methods

### Animals, Metabolic dysfunction-associated steatohepatitis-HCC mouse model and ethics statement

Adult C57BL/6J mice (8–16 weeks old) were obtained from the animal facility of the *Centro Regional de Estudios Avanzados para la Vida (CREAV)* at the Universidad de Concepción. The Metabolic dysfunction-associated steatohepatitis–HCC mouse model was generated in male mice by administering a single subcutaneous injection of 200 μg streptozotocin in 0.1 M citrate buffer (pH 4.5) at postnatal day 2, followed by feeding with a high-fat diet (32% fat, ad libitum) starting at 4 weeks of age(Fujii et al., 2013). Under this regimen, hepatocellular carcinoma developed by 16 weeks. Animals were housed under strict pathogen-free conditions and provided ad libitum access to food and water. All experimental procedures were approved by the Ethics and Biosecurity Committee of the Vice-Rectory for Research and Development at the Universidad de Concepción (ethics approval CICUA 2023-2025). The study adhered to ARRIVE guidelines, and all methods were conducted in accordance with institutional and national regulations.

### *Drosophila* brain collection and immunostaining

*Drosophila* third instar larval brains were dissected in phosphate-buffered saline (PBS), fixed in 4% paraformaldehyde (PFA) in PBS for 20 min. Samples were washed 3 times in PBT (1X PBS with 0.3% Triton X-100) and blocked in 1% bovine serum albumin (BSA)/PBT. Brains were then incubated overnight at 4 °C with mouse anti-Repo (DSHB clone 8D12, 1:20), a glia-specific nuclear marker. Samples were washed with PBT and incubated with anti-mouse Alexa Fluor 647 at 4°C overnight (ON). They were washed with PBT and mounted in Fluoroshield (SIGMA F6182).

### Mouse liver sample collection and immunostaining

Livers were fixed by intracardiac perfusion with 4% paraformaldehyde (PFA) in PBS supplemented with 0.1% Tween-20, followed by ON post-fixation in the same solution at room temperature. Liver samples were then sectioned into 100–200 μm slices using a vibratome. Tissues were stained with Alexa Fluor 488 Phalloidin (LIFE technologies, cat A12379, 1/1000) to label cell borders and DAPI (LIFE technologies, cat D1306, 1 µg/ml) to visualize nuclei. Finally, optical clearing was performed using a modified SeeDB protocol(Ke et al., 2013), as previously described(Morales-Navarrete et al., 2019).

### Liver and *Drosophila* sample imaging

Liver samples were imaged at a voxel size of 0.3 μm using an inverted multiphoton laser-scanning microscope (Zeiss LSM 780) equipped with a 40X, 1.2 NA multi-immersion objective (Zeiss). DAPI was excited at 780 nm with a Chameleon Ti–Sapphire two-photon laser, while Alexa Fluor 488 was excited using a 488 nm laser line and detected with gallium arsenide phosphide (GaAsP) detectors. For adult mouse livers, a single CV–PV axis from a randomly selected lobe was imaged, requiring a 2×1 tile. For HCC samples, imaging was performed across the tumor–adjacent tissue interface, which required a 3×2 tile to capture both regions.

*Drosophila* brain samples were imaged with an inverted multiphoton laser-scanning microscope (Zeiss LSM 780) equipped with a Plan-Apochromat 63x/1,4 Oil DIC (Zeiss), excited at 633 nm laser and acquiring a voxel size of 0.15×0.15×0.3 μm.

### Image pre-processing of Mouse and *Drosophila* images

To correct the irregular borders observed in the phalloidin and DAPI channels—arising from acquisition at maximum scanning speed without line-by-line averaging—we implemented a custom border-correction step based on the TV-L1 optical flow algorithm (skimage.registration.optical_flow_tvl1)(Besnerais and Champagnat, 2005; Plyer et al., 2016). For each 3D stack (ZYX), we exploited the fact that alternating Y-columns were shifted in opposite directions along the fast-scan axis: the volume was split into even and odd columns, each sub-volume was resampled back to the original shape, and a dense optical flow field was estimated between them. The even and odd sub-volumes were then warped symmetrically towards the center (± flow/2), and their interleaved columns were recombined to obtain a single stack with aligned borders. To reconstruct the full CV–PV axis or the tumor– adjacent tissue interface, image tiles were stitched using the stitching plug-in in Fiji(Preibisch et al., 2009). DAPI and phalloidin signal intensities were then normalized along the Z-axis using the 16-bit cumulative histograms (65,536 bins), rescaling the 10th–99.99th and corrected using the *Bleach Correction* plugin from Fiji(Miura, 2020) with the Histogram Matching method, and deconvolution was performed using the SelfNet network(Ning et al., 2023).

### Manual annotation of nuclei in mouse and *Drosophila* images

Nuclei were manually annotated in Z-stacks obtained from livers of 3 adult mice (8–10 weeks old), 4 adult mice (16 weeks old), 4 adult mice with HCC, and from 3 *Drosophila* brains, using Napari (https://zenodo.org/records/18374344). Annotations were performed in 2D and subsequently interpolated in 3D using the napari-nD-annotator plugin with the RPSV interpolation method, as described in the corresponding GitHub repository (https://github.com/bauerdavid/napari-nD-annotator). The total number of annotated nuclei was 2,873 nuclei across adult mouse livers, 2,651 nuclei across HCC livers, and 702 nuclei across *Drosophila* brains. All manual annotations are publicly available via Zenodo(https://doi.org/10.5281/zenodo.18517324).

### 3D Segmentation and morphological analysis of HCC liver tissue

Nuclear and cellular morphological analyses were performed using MotionTracking (http://motiontracking.mpi-cbg.de)(Morales-Navarrete et al., 2015). Triangulated meshes were generated from the segmented nuclei produced by NucVerse3D. Hepatocyte surfaces were then reconstructed using an active mesh expansion approach, in which each nuclear mesh was iteratively expanded until reaching the cell-border marker. For a detailed description of this procedure, see(Morales-Navarrete et al., 2015).

### 3D network architecture and training (NucVerse3D)

A central challenge in 3D instance segmentation is that instance identities cannot be learned directly from voxel wise integer labels. Unlike semantic masks, instance labels are inherently arbitrary: the same nucleus may carry a different integer index across images, and the ordering of labels has no physical or biological meaning. Consequently, a neural network cannot be trained to regress these labels directly. To overcome this limitation, instance segmentation models require a geometry-aware, label-invariant representation that (i) can be predicted consistently across images and (ii) preserves sufficient information to reconstruct the original instance masks.

In NucVerse3D, we address this challenge by transforming each ground-truth nucleus into a continuous and reversible 3D gradient-field representation. For each nucleus, we compute its geometric centroid and, for every voxel within the corresponding mask, derive a displacement vector encoding the Euclidean offsets in the x, y, and z directions relative to that centroid. These displacement vectors are normalized by their magnitude, producing a unit directional field in which each voxel stores the direction pointing from the centroid to that voxel. The result is a smooth, modality-agnostic 3D vector field whose components lie in the range [−1,1][-1, 1][−1,1], capturing the local geometry and orientation of each nucleus without relying on discrete instance identifiers.

During inference, the network jointly predicts (i) a binary foreground probability map and (ii) the three components of the 3D gradient field. Instance reconstruction proceeds through a gradient-descent–based grouping procedure. Starting from all voxels classified as foreground, each voxel is iteratively displaced against its predicted gradient direction, effectively following the inverse of the outward flow, until convergence. Voxels whose trajectories converge to a similar spatial location are considered to share a common attractor, representing an estimated nucleus centroid. These attractor points are subsequently clustered to produce a set of centroid hypotheses, each of which is assigned a unique instance label.

Finally, instance masks are recovered by assigning each foreground voxel the label associated with the attractor to which its gradient-descent path converged. This yields a complete set of 3D instance masks, fully reconstructed from the predicted auxiliary representation. Importantly, the entire process is reversible, differentiable, and label-invariant, enabling the network to learn complex nuclear shapes and boundaries in a geometrically meaningful space without the ambiguity inherent in integer instance labels.

### Preprocessing (NucVerse 3D)

All volumetric images were preprocessed to correct uneven illumination, enhance image quality, and normalize intensity values prior to training.

#### i Correction of uneven illumination

Uneven illumination was corrected in a model-free manner using a ratiometric approach. Given a raw image volume *I*_*raw*_, a low-frequency background illumination field *I*_*bg*_ was estimated by first downsampling the image by a factor *ds* = *8*, applying isotropic Gaussian smoothing with standard deviation σ = *5*, and resizing the result back to the original image resolution using linear interpolation. Background values below a threshold of τ = *0*.*01* were clipped to prevent numerical instabilities. The corrected image *I*_*correct*_ as

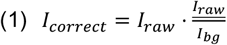

where *I*<u>_*raw*_</u> denotes the mean intensity of the raw image.

#### ii Unsupervised deconvolution

Following illumination correction, blind deconvolution was performed using the unsupervised Wiener–Hunt approach implemented in *scikit-image*(Orieux et al., 2010). An initial point-spread function (PSF) was defined as a sum-normalized 2D Gaussian kernel of size *15* × *15* pixels with standard deviation σ_+,-_ = *1*. Deconvolution was applied independently to each axial slice of the 3D volume.

#### iii Intensity normalization

Finally, contrast-enhancing intensity normalization was applied using percentile-based rescaling. Image intensities were mapped to the [0,1] range by setting the lower and upper bounds to the 2nd and 99.8th percentiles of the intensity distribution, respectively. This normalization suppresses low-level background signals and clips sporadic high-intensity outliers while preserving relative contrast between nuclear structures.

### Nuclear size normalization (scaling)

To reduce inter-dataset variability in nuclear size and to stabilize training across modalities, we implemented an isotropic nuclear scaling step prior to patch extraction. For each training volume, we first estimated the mean nuclear radius from the available ground-truth annotations. The raw 3D image was then resampled isotropically such that the average nuclear radius was mapped to approximately 10 µm. This global scaling was applied uniformly to all images in the set, preserving relative nuclear morphology while harmonizing object size across datasets.

### Data Augmentation (NucVerse 3D)

To reduce overfitting and improve robustness, 3D data augmentation was applied on-the-fly during training and jointly to images and labels. Geometric augmentations included random flips along the three spatial axes, axis transpositions, and random 3D rotations combined with anisotropic scaling. Intensity-based augmentations comprised random brightness variations using a power-law transformation, Gaussian blurring, and Poisson noise. In addition, elastic deformations were applied to introduce non-linear spatial variability. At least 25% of the training samples were left unaugmented to preserve the original data distribution.

### Other models

#### i StarDist 3D

(Weigert and Schmidt, 2022): The same processed datasets used to train NucVerse3D were also used for training StarDist 3D. From these datasets, 3D patches of size 64 × 96 × 96 voxels were extracted, and the same data augmentation strategy was applied. The model was trained for 200 epochs with a batch size of 2 and an initial learning rate of 1 × 10^−3^.

#### ii CellPose 3.0

(Stringer and Pachitariu, 2025): The Cellpose model was trained using the same datasets as NucVerse3D. For each dataset, the original 3D volumes were first preprocessed following the same pipeline described previously (correction of uneven illumination, unsupervised deconvolution and intensity normalization), and then converted into a list of 2D slices for training. Slices whose corresponding masks contained fewer than 20 labeled pixels were removed before training. Since normalization was already applied during preprocessing, Cellpose internal normalization was disabled. The model was trained for 200 epochs with a learning rate of 1 × 10^−3^, using a 90/10 train–validation split.

### Segmentation benchmark datasets

Seven volumetric microscopy datasets of cellular nuclei were used to evaluate the generalization of NucVerse3D across species, tissues, and imaging modalities. Three datasets were generated in this study, including (1) two-photon microscopy Z-stacks of adult mouse liver tissue control, (2) two-photon microscopy Z-stacks of HCC, and (3) confocal microscopy volumes of *Drosophila melanogaster* brain regions stained for glial nuclei. All manual annotations are publicly available via Zenodo(https://doi.org/10.5281/zenodo.18517324). The remaining datasets were obtained from previously published studies.

#### i Mouse Liver Tissue Dataset

The Mouse liver dataset comprises two-photon microscopy stacks of adult mouse liver tissue containing mono- and binucleated hepatocytes, acquired at a voxel size of 0.3 × 0.3 × 0.3 µm. The training subset consists of 6 volumes with 2,016 annotated nuclei, while the test subset includes 3 volumes containing 857 nuclei.

#### ii *Drosophila* Brain Dataset

This dataset includes confocal microscopy volumes of *Drosophila melanogaster* brain regions stained for glial cells, acquired with a voxel size of 0.15 × 0.15 × 0.3 µm. The training set comprises 3 volumes with 399 annotated nuclei, while the test set contains 2 volumes with 303 nuclei.

#### iii Mouse Liver HCC Dataset

The Mouse Liver HCC dataset comprises two-photon microscopy volumes of HCC tissue, acquired at a voxel size of 0.3 × 0.3 × 0.3 µm. The training subset consists of 8 volumes with 1,750 annotated nuclei (218.75 ± 46.08 nuclei per volume), while the test subset includes 4 volumes containing 901 nuclei (225.25 ± 28.86 nuclei per volume).

#### iv C. elegans 3D Fluorescence Microscopy Nuclei Dataset

(Hirsch and Kainmueller, 2020; Long et al., 2009): This dataset consists of confocal microscopy volumes of *Caenorhabditis elegans* worms at the L1 developmental stage, acquired with a voxel size of 0.116 × 0.116 × 0.122 µm. The dataset is split into training, validation, and test subsets. The training set contains 21 volumes with a total of 11,387 annotated nuclei and the test set comprises 10 volumes containing 3,761 nuclei. All data are publicly available through Zenodo (https://zenodo.org/records/5942575).

#### V Whole Mouse Brain Nuclei Dataset

(Achard et al., 2025): The Whole Mouse Brain dataset was acquired using light-sheet mesoSPIM imaging of an entire mouse brain at a coarse voxel size of 5.26 × 5.26 × 5 µm. The training split includes 5 volumes with a total of 2,082 nuclei, while the test split consists of 2 volumes containing 559 nuclei. The dataset is publicly available through Zenodo (https://zenodo.org/records/11095111).

#### vi-vii NucMM Challenge Datasets

(Lin et al., 2021): The remaining datasets were obtained from the NucMM Challenge and include micro-computed tomography (microCT) volumes of mouse visual cortex acquired at a voxel size of 0.72 × 0.72 × 0.72 µm, as well as scanning electron microscopy (SEM) volumes of zebrafish brain acquired at submicron resolution (0.48 × 0.51 × 0.51 µm). For the mouse visual cortex dataset, the training set contains 4 volumes with 572 annotated nuclei, and the test set comprises 4 volumes containing 664 nuclei. For the zebrafish brain dataset, the training set contains 27 volumes with 7,271 annotated nuclei, and the test set comprises 27 volumes containing 8,044 nuclei. These datasets are publicly available through the NucMM Challenge platform (https://nucmm.grand-challenge.org/NucMM/).

### Quantification of segmentation quality

For each image, segmentation quality was evaluated at the instance level by computing an optimal matching between predicted and reference cell instances. Given a set of M predicted instances and N ground-truth instances, predicted nuclei are matched to reference nuclei using a one-to-one correspondence based on spatial overlap. For each predicted instance, the intersection-over-union (IoU) metric was computed with candidate reference instances, and only pairs with non-zero overlap were considered valid matches. Unmatched predictions were counted as false positives (FP), while reference instances with no corresponding match were counted as false negatives (FN).

#### i Intersection-over-Union (IoU)

The intersection-over-union (IoU), also known as the Jaccard index, is defined as the ratio between the number of voxels in the intersection and the number of voxels in the union of two binary instance masks A and B. IoU values range from 0 to 1, with higher values indicating greater spatial overlap. IoU was used as the primary criterion to determine whether a predicted instance is considered a correct detection according to a predefined threshold.

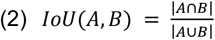

#### ii Precision, Recall and F1-Score

Based on the instance matching defined by the IoU criterion, segmentation performance was further quantified using precision, recall, and the F1-score. Precision measures the proportion of predicted instances that are correctly matched to a ground-truth object, reflecting the reliability of the detected nuclei. Recall quantified the fraction of ground-truth instances that are successfully recovered by the model, capturing its completeness. The F1-score was computed as the harmonic mean of precision and recall and provides a single summary metric that balances over-segmentation and under-segmentation effects, making it particularly suitable for evaluating instance-level cell segmentation performance.

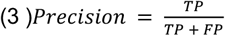

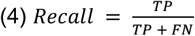

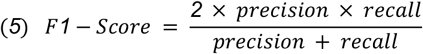

#### iii Average Precision (AP) curve

In addition to single-threshold metrics, segmentation performance was evaluated using the average precision (AP). For each image, predicted instances were evaluated across multiple IoU thresholds, where lower thresholds require less spatial overlap for a valid match. At each IoU threshold, valid matches defined the true positives, while unmatched predictions and reference instances defined false positives and false negatives, respectively. The AP at a given threshold was computed as

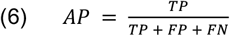

The AP curve for a dataset was obtained by averaging the AP values across all images and across a set of uniformly sampled IoU thresholds in the range [0.5,1.0].

### Quantification of nuclear content–size coupling and abnormality scoring

#### i Nuclear content–volume scaling model

To quantify deviations between nuclear DNA content and nuclear size at the single-nucleus level, we modeled the expected relationship between nuclear volume and DAPI intensity using a log–log linear framework. For each segmented nucleus *i*, nuclear volume *Vi* and integrated DAPI intensity *Ii* were computed from the 3D nuclear masks generated by NucVerse3D. To compensate for variation in imaging condition, the DAPI intensities were normalised by dividing them by the median integrated DAPI intensity of the non-parenchymal nuclei in the corresponding samples as they correspond to diploid nuclei. Both quantities were log-transformed using base 2 to ensure interpretability of scaling coefficients and variance stabilization, as shown in eq. 7:

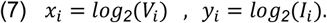

A linear regression model was fitted using nuclei from control tissue only:

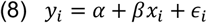

where α and β represent the intercept and scaling exponent, respectively.

This control-derived model defines the expected coupling between nuclear size and DNA content under physiological conditions and was subsequently applied unchanged to all experimental conditions.

#### ii Residual computation and normalization

For each nucleus, the signed residual was computed as:

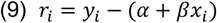

which captures whether a nucleus contains more (*r*_*i*_ > *0*) or less (*r*_*j*_ < *0*) DNA than expected for its size.

To enable cross-sample and cross-condition comparability, residuals were normalized by the standard deviation of residuals observed in control tissue:

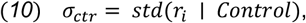

yielding two complementary scores:

- Signed Nuclear Decoupling Score (sNDS):

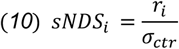
- Unsigned Nuclear Decoupling Score (NDS):

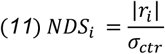

By construction, whereas sNDS preserves the direction of decoupling (DNA excess vs. deficit), NDS quantifies the magnitude of abnormality irrespective of sign.

#### iii Definition of nuclear abnormality classes

Nuclei were classified into discrete abnormality categories based on their NDS values, using thresholds defined relative to control variability. The primary binary definition of abnormality used throughout the study was NDS>2, corresponding to deviations exceeding two standard deviations from the control distribution. This threshold captures nuclei unlikely to arise from physiological variability and is robust to moderate deviations from normality in the residual distribution.

#### iv Sample-level aggregation and robustness

To avoid bias from unequal nucleus counts across samples, all downstream statistical analyses were performed at the sample level. For each sample, we computed the fraction of nuclei classified as abnormal (NDS>2). This aggregation ensures that each biological replicate contributes equally to group-level comparisons.

Robustness of the decoupling framework was assessed by inspecting residual distributions, sNDS density plots, and NDS histograms across conditions, confirming that control nuclei were centered near zero with unit variance, while adjacent and tumor tissues exhibited broader, shifted distributions.

#### v Statistical analysis

Differences in the fraction of abnormal nuclei between conditions were evaluated using one-sided Mann–Whitney U tests, reflecting the a priori hypothesis of increased nuclear abnormality from control to adjacent tissue and from adjacent tissue to tumor. Specifically, the following directional comparisons were tested: Control < Adjacent, Control < Tumor, and Adjacent < Tumor.

To correct for multiple testing, p-values were adjusted using the Benjamini–Hochberg false discovery rate (FDR) procedure. Statistical significance was reported using standard star notation (*p* < 0.05, p < 0.01, *p* < 0.001). All analyses were performed using SciPy and statsmodels in Python.

## Research data

The source code for training and predicting nuclei segmentation using NucVerse 3D is available from https://github.com/Segovia-lab/3D-Nuclei-segmentation.git

## Data statement

All data generated in this study are publicly available through Zenodo (https://doi.org/10.5281/zenodo.18517324), including: For the newly released datasets, (1) raw volumetric datasets, (2) manual annotations, (3) preprocessed patches used for model training, (4) trained model weights, and (5) segmentation outputs. For external benchmark datasets, only the (6) preprocessed patches used for model training, (7) trained model weights, and (8) segmentation outputs are provided, while the raw data are available from the original sources.

## Acknowledgements

We thank the Corporación Ecuatoriana para el Desarrollo de la Investigación y la Academia (CEDIA) for providing access to its high-performance computing resources and technical support, which enabled the computational work for this study. We also thank the contributors of the open-source tools utilized in this work.

## Author Contributions

**J.V**.: Conceptualization, Formal analysis, Investigation, Methodology, Software, Writing – Original draft preparation. **C.P.-G**.: Data curation, Formal analysis, Investigation, Methodology. **R.V**.: Formal analysis, Investigation, Methodology, Software. **D.M**.: Data curation, Formal analysis, Investigation. **D.B**.: Data curation. **E.G.C**.: Data curation, Methodology, Writing – Reviewing and Editing, Funding acquisition. **P.G**.: Conceptualization, Writing – Reviewing and Editing, Funding acquisition. **F.S.-M**.: Conceptualization, Formal analysis, Writing – Original draft preparation, Funding acquisition, Supervision. **H.M.-N**.: Conceptualization, Formal analysis, Investigation, Software, Writing – Original draft preparation, Supervision.

## Funding sources

The authors acknowledge the funding support from ANID, and Instituto Milenio Centro Interdisciplinario de Neurociencia de Valparaíso (CINV), under grant numbers ANID Fondecyt regular 1251048, and ICN2025_026 CINV to FS-M. FONDECYT Iniciación en Investigación 11230539 to E.G.C. ANID AC3E CIA250006 and CENIA FB210017 to P.G.

## Competing Interests Statement

The authors declare that they do not have competing interests.

## Declaration of generative AI and AI-assisted technologies in the manuscript preparation process

During the preparation of this work the author(s) used ChatGPT 5.2 in order to rephrase some sections of the manuscript and correct grammatical mistakes. After using this tool/service, the author(s) reviewed and edited the content as needed and take(s) full responsibility for the content of the published article.

## Figure and table legends

**Supplementary Figure 1:**
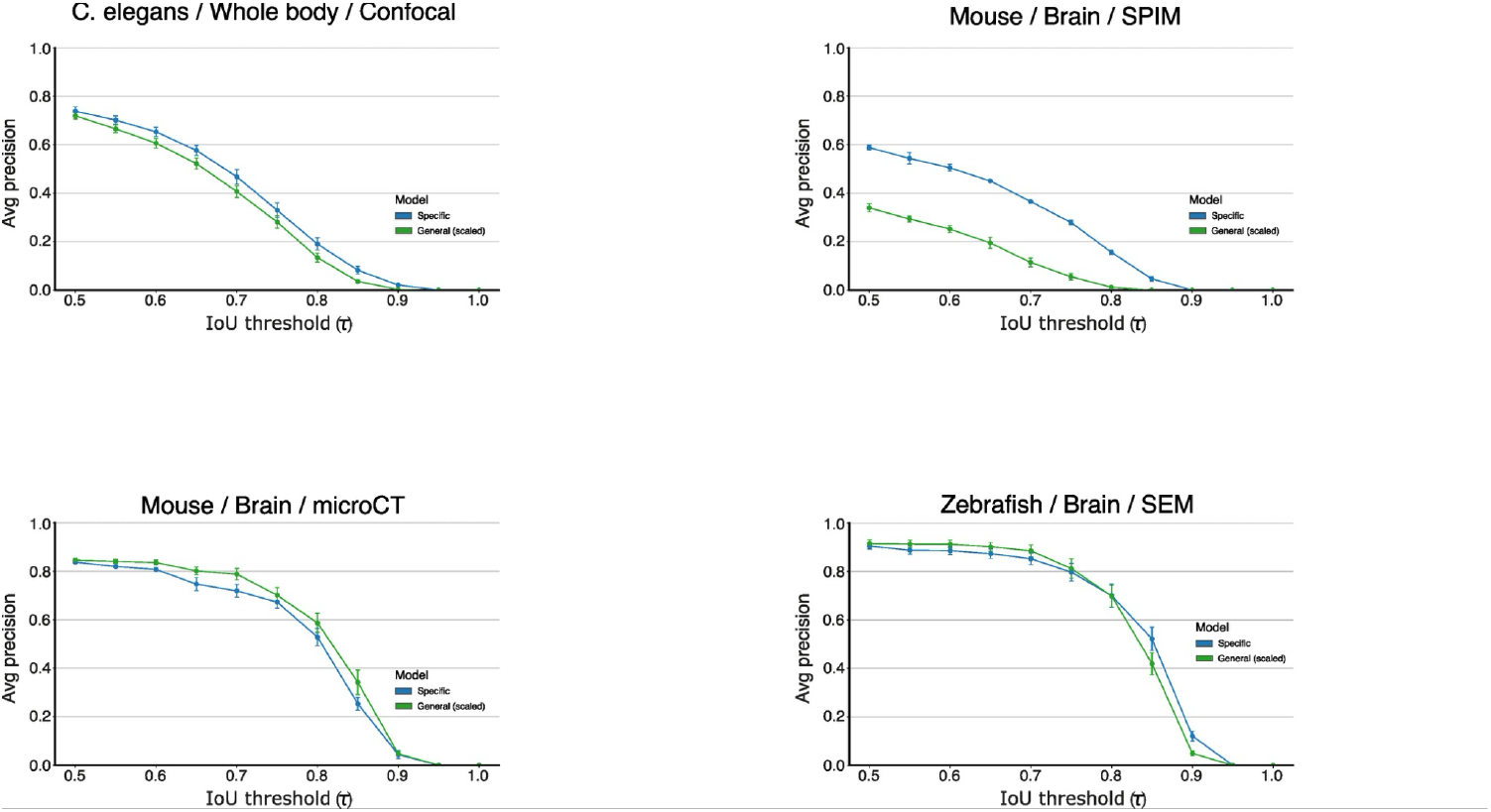
Performance of generalized and dataset-specific models across imaging modalities. Average precision (AP) versus intersection-over-union (IoU) threshold (τ) for dataset-specific and generalized NucVerse3D models across four volumetric microscopy datasets: *C. elegans* whole-body confocal, mouse brain SPIM, mouse brain microCT, and zebrafish brain SEM. The generalized model shows performance comparable to dataset-specific models across imaging modalities and species. Error bars represent mean ± standard error of mean across test volumes (*C. elegans* confocal, n = 7; mouse brain SPIM, n = 2; mouse brain microCT, n = 4; zebrafish brain SEM, n = 27; Drosophila brain confocal, n = 2; mouse liver 2-photon, n = 3).

**Supplementary Figure 2:**
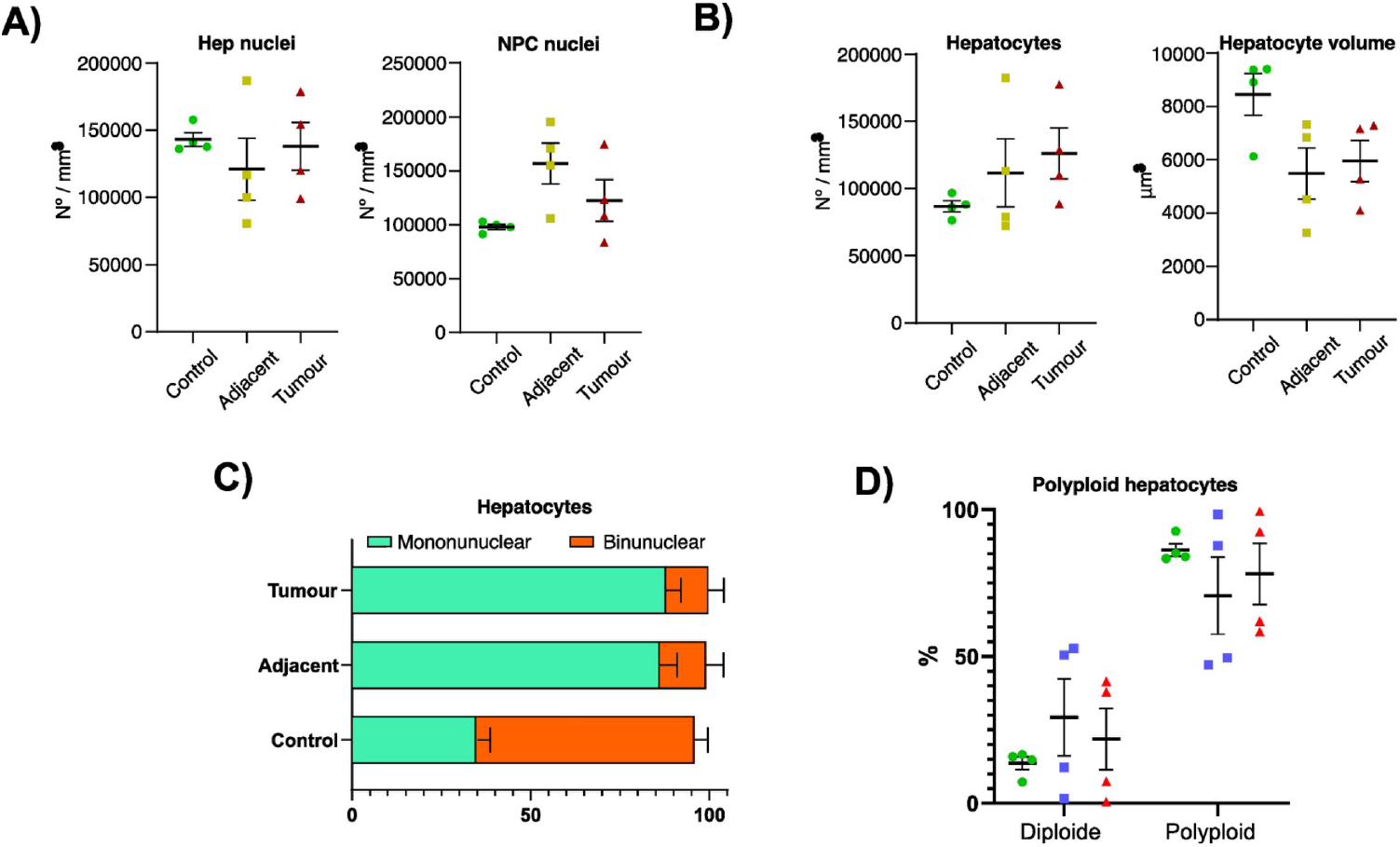
Nuclear- and cell-level quantification of hepatocyte organization and ploidy across control, adjacent, and tumor liver tissue. Quantitative analysis of nuclear- and cell-level features extracted from 3D segmentations of liver tissue under control, and HCC conditions (adjacent and tumour). **(A)** Density of hepatocyte (Hep) nuclei and non-parenchymal cell (NPC) nuclei, normalized by tissue volume. **(B)** Density of hepatocytes per unit volume (left) and mean hepatocyte volume (right). **(C)** Relative proportions of mononuclear and binuclear hepatocytes across conditions. **(D)** Fraction of diploid and polyploid hepatocytes per condition. Error bars represent mean ± standard error of mean across control livers (n = 4), and HCC livers (n = 4).

**Supplementary Figure 3:**
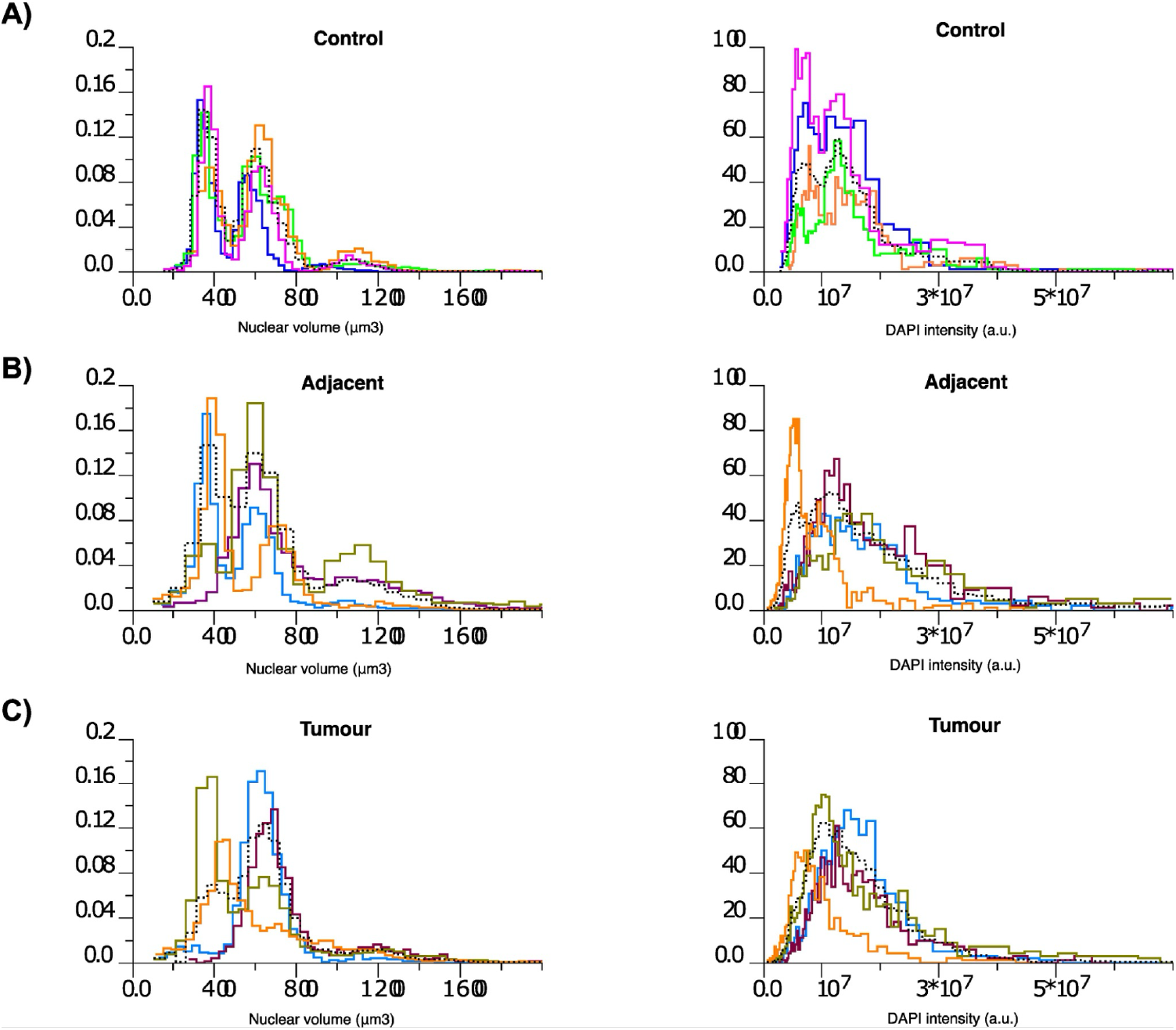
Distributions of nuclear volume and DNA content across control, adjacent, and tumor liver tissue. Distributions of nuclear volume (left panels) and integrated DAPI intensity (right panels) for individual nuclei across liver tissue conditions: **(A)** control, **(B)** adjacent, and **(C)** tumor. Colored curves represent individual biological samples, while the black dotted line indicates the aggregate distribution across all samples within each condition. Analyses were performed on 4 biological samples per condition.

**Supplementary Figure 4:**
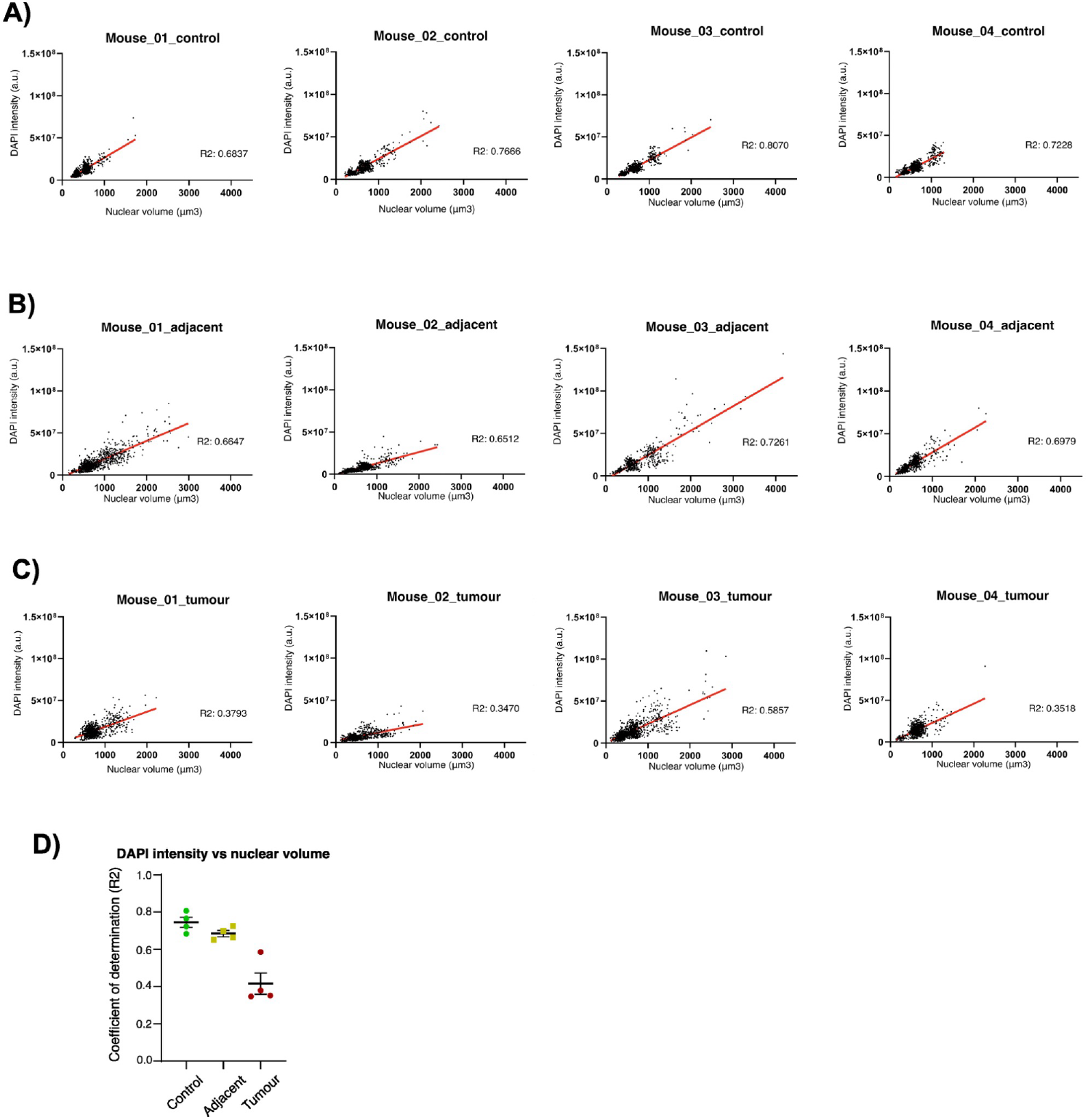
Sample-level coupling between nuclear volume and DNA content in control, adjacent, and tumor liver tissue. Scatter plots showing the relationship between nuclear volume and integrated DAPI intensity for individual nuclei, displayed separately for each biological sample. Panels correspond to **(A)** control, **(B)** adjacent, and **(C)** tumor liver tissue. Each dot corresponds to a single nucleus, and the red line indicates the fitted linear regression for each sample. The coefficient of determination (R^2^) is reported for each fit. **(D)** Sample-level coefficient of determination (R^2^) quantifying the coupling between integrated DAPI intensity and nuclear volume.

**Supplementary Figure 5:**
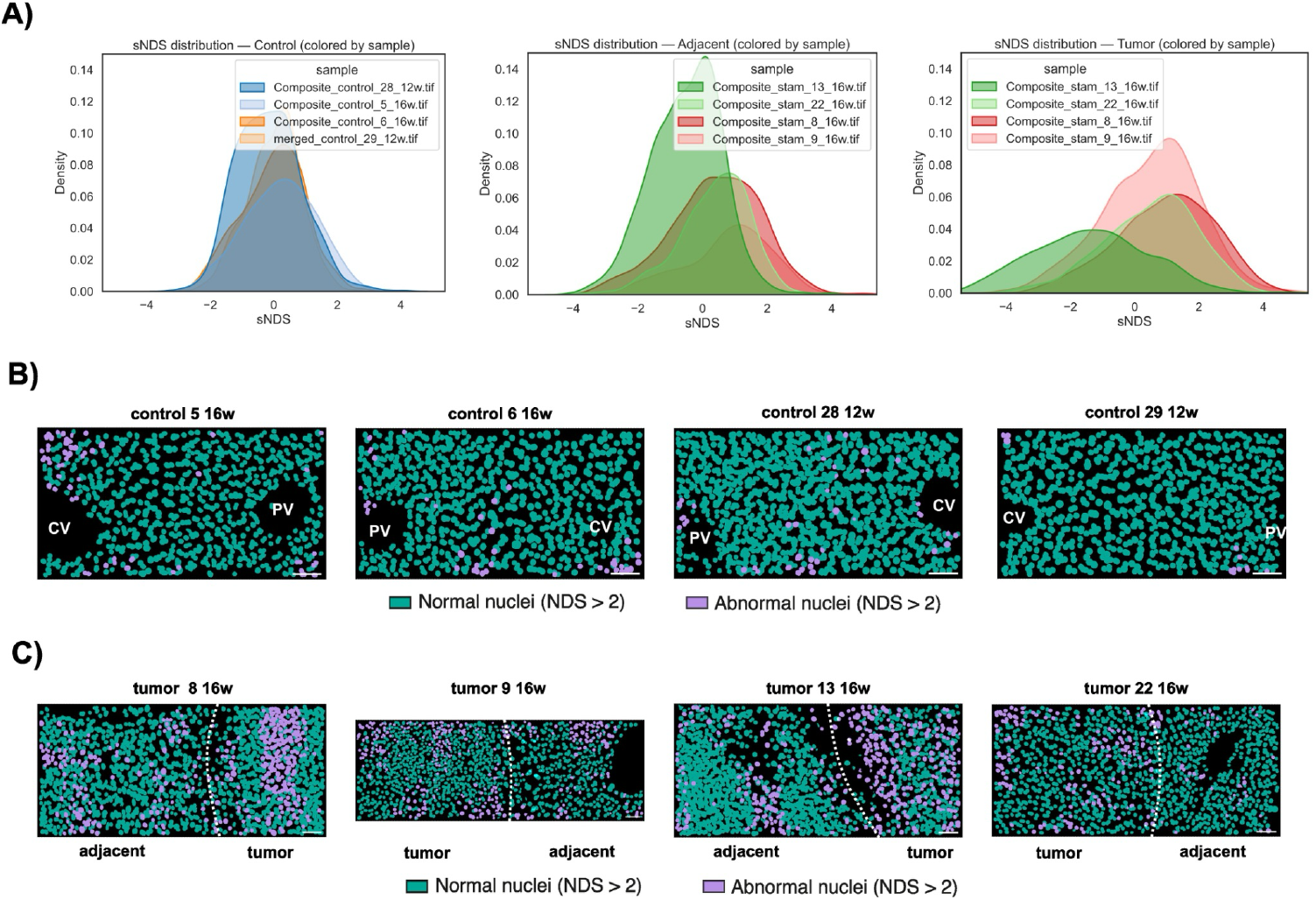
Sample-level distributions and spatial mapping of standardized nuclear deviation scores (sNDS) in control, adjacent, and tumor liver tissue. **(A)** NDS distributions for individual samples, shown separately for control, adjacent, and tumor conditions. **(B)** Spatial maps of nuclei classified as normal or abnormal based on sNDS in control and HCC **(C)** liver samples. Each panel shows a maximum-intensity projection over a 50 μm depth, with central vein (CV) and portal vein (PV) regions indicated. Scale bars 50 µm.

**Supplementary Table 1:** Summary of datasets used for training and evaluation

**Supplementary Movie 1: 3D instance segmentation workflow with NucVerse3D**. 3D rendering of DAPI-stained nuclei in a mouse liver sample, followed by the corresponding 3D nuclear mask with detected centroids, and the final 3D instance segmentation.

**Supplementary Movie 2: 3D nuclear instance segmentation in *C. elegans***. The movie shows the raw confocal Z-stack from a *C. elegans* dataset as sequential optical sections along the axial (Z) dimension, followed by nuclear predictions generated by NucVerse3D, displayed as nuclear outlines and the corresponding 3D nuclear instance masks.

**Supplementary Movie 3: 3D nuclear instance segmentation in mouse brain (SPIM)**. The movie shows the raw light-sheet Z-stack from a mouse brain dataset as sequential optical sections along the axial (Z) dimension, followed by nuclear predictions generated by NucVerse3D, displayed as nuclear outlines and the corresponding 3D nuclear instance masks.

**Supplementary Movie 4: 3D nuclear instance segmentation in mouse visual cortex (microCT):** The movie shows the raw microCT volume from a mouse visual cortex dataset as sequential axial sections, followed by nuclear predictions generated by NucVerse3D, displayed as nuclear outlines and the corresponding 3D nuclear instance masks.

**Supplementary Movie 5: 3D nuclear instance segmentation in zebrafish brain (SEM):** The movie shows the raw SEM volume from a zebrafish brain dataset as sequential axial sections, followed by nuclear predictions generated by NucVerse3D, displayed as nuclear outlines and the corresponding 3D nuclear instance masks.

**Supplementary Movie 6: 3D nuclear instance segmentation in *Drosophila melanogaster* brain:** The movie shows the raw confocal Z-stack of glial cells from a *Drosophila melanogaster* brain dataset as sequential optical sections along the axial (Z) dimension, followed by nuclear predictions generated by NucVerse3D, displayed as nuclear outlines and the corresponding 3D nuclear instance masks.

**Supplementary Movie 7: 3D nuclear instance segmentation in adult mouse liver:** The movie shows the raw two-photon Z-stack from an adult mouse liver dataset as sequential optical sections along the axial (Z) dimension, followed by nuclear predictions generated by NucVerse3D, displayed as nuclear outlines and the corresponding 3D nuclear instance masks.

**Supplementary Movie 8: Comparison of dataset-specific and generalized models in *Drosophila melanogaster* brain:** The movie shows the raw confocal Z-stack from a *Drosophila melanogaster* brain dataset as sequential optical sections along the axial (Z) dimension, followed by nuclear outlines predicted by NucVerse3D using dataset-specific and generalized models.

**Supplementary Movie 9: Comparison of dataset-specific and generalized models in adult mouse liver:** The movie shows the raw two-photon Z-stack images from an adult mouse liver dataset as sequential optical sections along the axial (Z) dimension, followed by nuclear outlines predicted by NucVerse3D using dataset-specific and generalized models.

## Notes

### Competing Interest Statement

The authors have declared no competing interest.

### Summary of Updates

The link to the Zenodo repository was coorected, now it is (https://doi.org/10.5281/zenodo.18517324)

