## Supplementary figures and images for "NucVerse3D: Generalizable 3D nuclear instance segmentation across heterogeneous microscopy modalities"

### Supp Fig 01

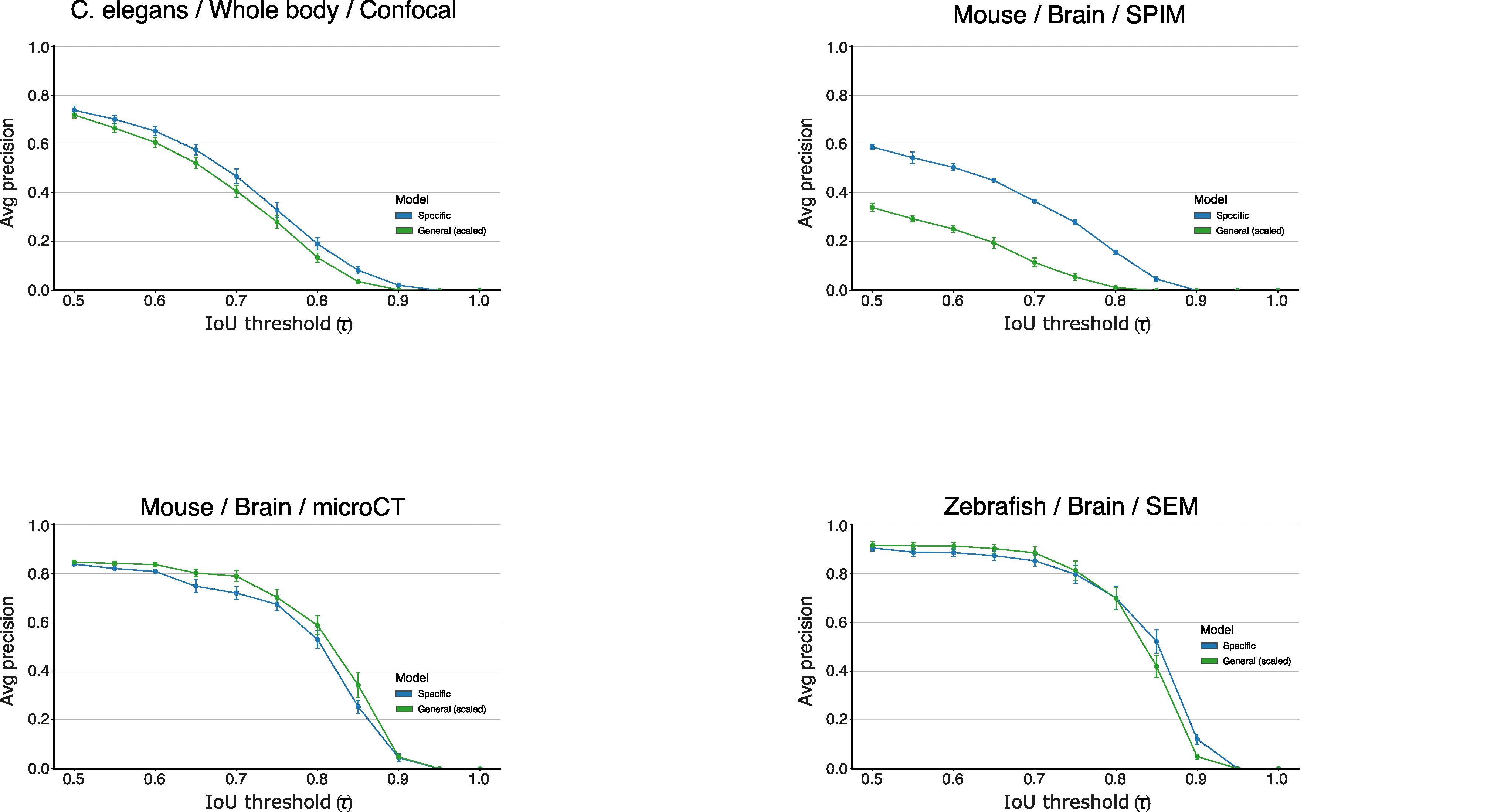

### Supp Fig 02

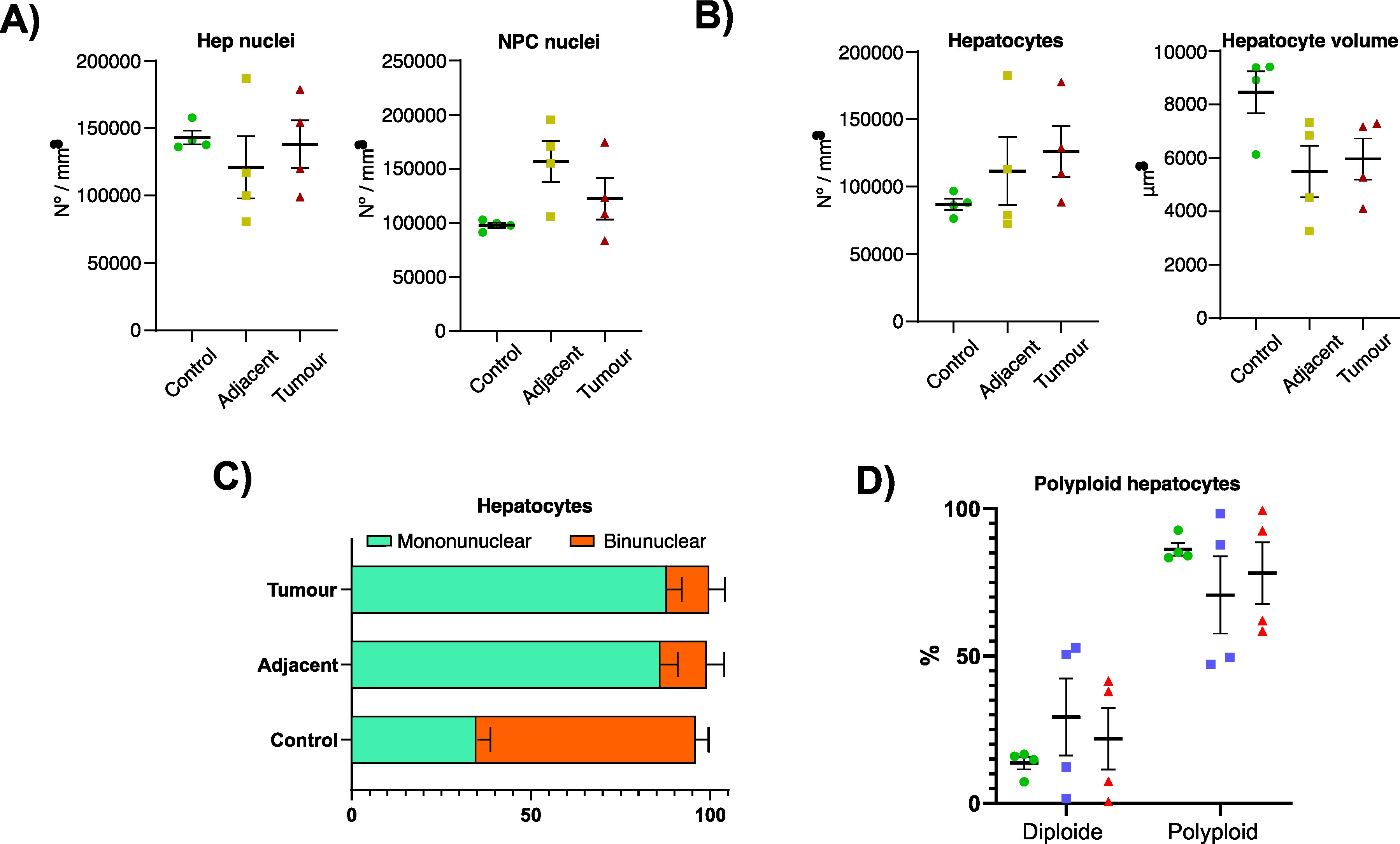

### Supp Fig 03

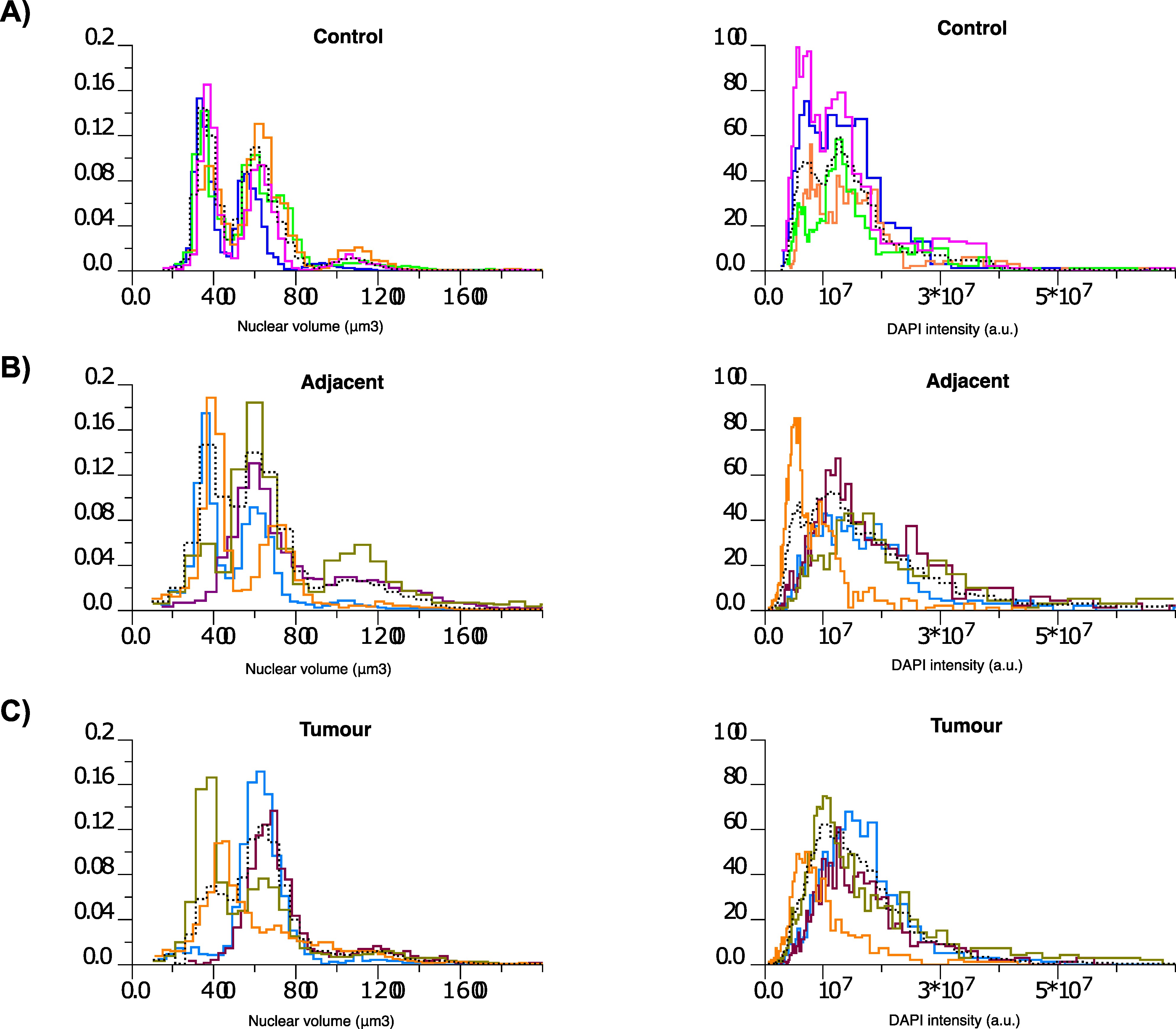

### Supp Fig 04

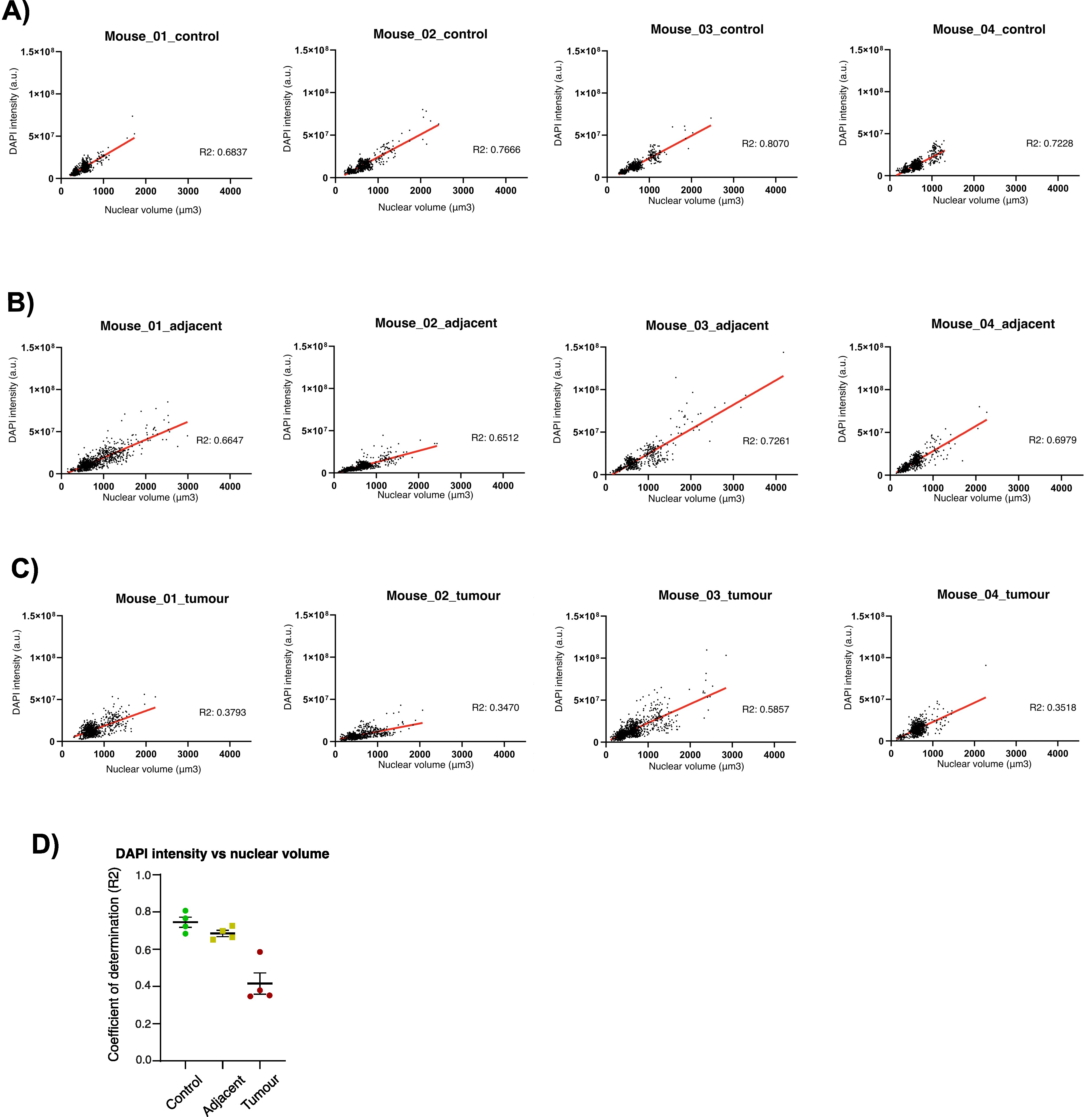

### Supp Fig 05

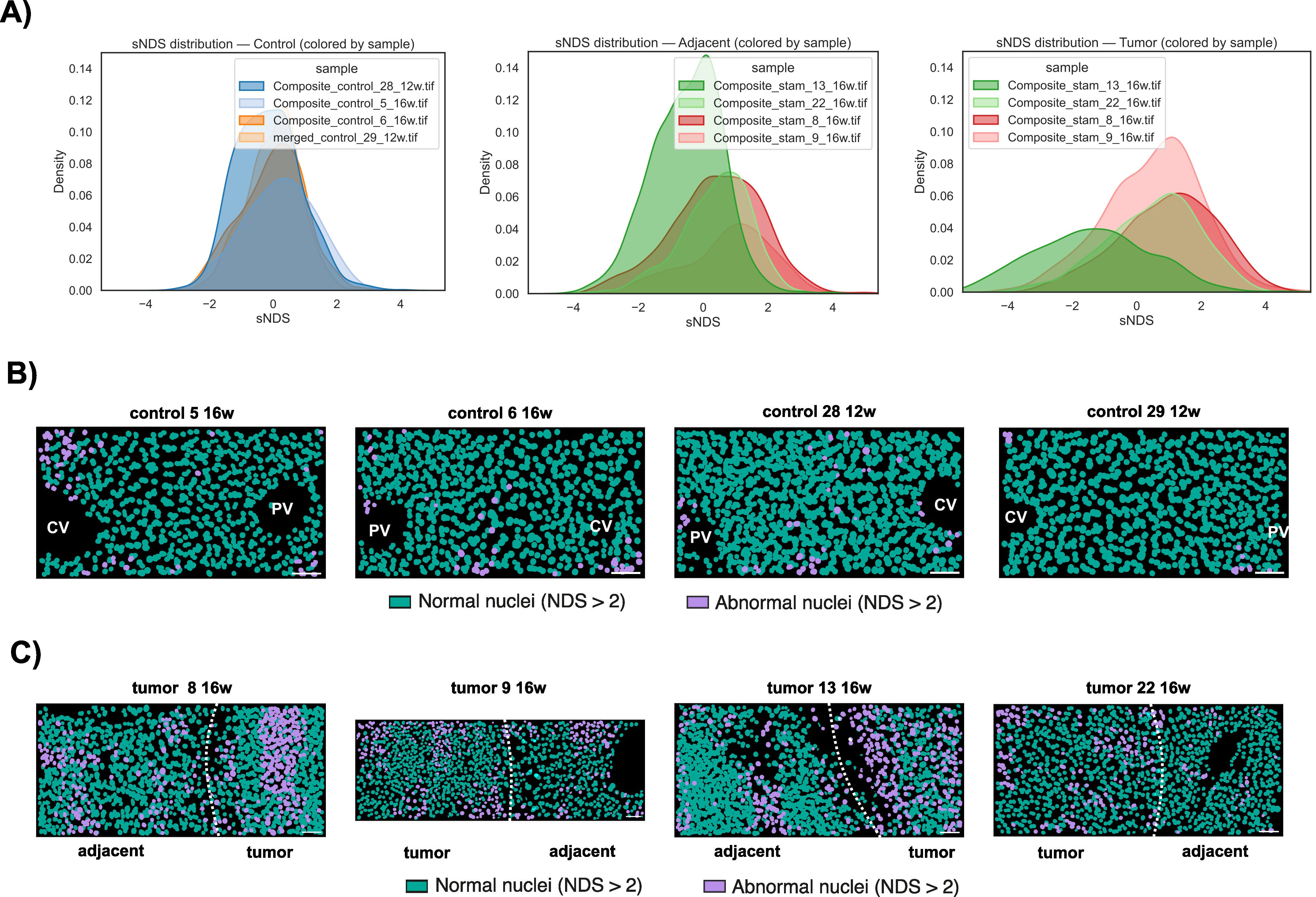
